# A resource to enable chemical biology and drug discovery of WDR Proteins

**DOI:** 10.1101/2024.03.03.583197

**Authors:** Suzanne Ackloo, Fengling Li, Magda Szewczyk, Almagul Seitova, Peter Loppnau, Hong Zeng, Jin Xu, Shabbir Ahmad, Yelena A Arnautova, AJ Baghaie, Serap Beldar, Albina Bolotokova, Paolo A Centrella, Irene Chau, Matthew A Clark, John W Cuozzo, Saba Dehghani-Tafti, Jeremy S Disch, Aiping Dong, Antoine Dumas, Jianwen A. Feng, Pegah Ghiabi, Elisa Gibson, Justin Gilmer, Brian Goldman, Stuart R Green, Marie-Aude Guié, John P Guilinger, Nathan Harms, Oleksandra Herasymenko, Scott Houliston, Ashley Hutchinson, Steven Kearnes, Anthony D Keefe, Serah W Kimani, Trevor Kramer, Maria Kutera, Haejin A Kwak, Cristina Lento, Yanjun Li, Jenny Liu, Joachim Loup, Raquel AC Machado, Christopher J Mulhern, Sumera Perveen, Germanna L Righetto, Patrick Riley, Suman Shrestha, Eric A Sigel, Madhushika Silva, Michael D. Sintchak, Belinda L Slakman, Rhys D Taylor, James Thompson, Wen Torng, Carl Underkoffler, Moritz von Rechenberg, Ian Watson, Derek J Wilson, Esther Wolf, Manisha Yadav, Aliakbar K Yazdi, Junyi Zhang, Ying Zhang, Vijayaratnam Santhakumar, Aled M Edwards, Dalia Barsyte-Lovejoy, Matthieu Schapira, Peter J Brown, Levon Halabelian, Cheryl H Arrowsmith

**Affiliations:** Structural Genomics Consortium, University of Toronto, 101 College St, Toronto, ON M5G 1L7, Canada; Google, 1600 Amphitheatre Parkway, Mountain View, CA 94043, USA; ZebiAI Inc., 100 Beaver St., Waltham MA 02435, USA; X-Chem Inc., 100 Beaver St., Waltham MA 02435, USA; Relay Therapeutics, 399 Binney St., Cambridge MA 02139, USA; Princess Margaret Cancer Centre, University of Toronto, Toronto, ON M5G 2M9, Canada; Civetta Therapeutics, 10 Wilson Rd, Cambridge, MA 02138, USA; Department of Chemistry, York University, Toronto, ON M3J 1P3, Canada; Department of Chemistry, University of Toronto, ON M5S 3H6, Canada; Department of Pharmacology and Toxicology, University of Toronto, Toronto, ON M5S 1A8, Canada

## Abstract

Protein class-focused drug discovery has a long and successful history in pharmaceutical research, yet most members of druggable protein families remain unliganded, often for practical reasons. Here we combined experiment and computation to enable discovery of ligands for WD40 repeat (WDR) proteins, one of the largest human protein families. This resource includes expression clones, purification protocols, and a comprehensive assessment of the druggability for hundreds of WDR proteins. We solved 21 high resolution crystal structures, and have made available a suite of biophysical, biochemical, and cellular assays to facilitate the discovery and characterization of small molecule ligands. To this end, we use the resource in a hit-finding pilot involving DNA-encoded library (DEL) selection followed by machine learning (ML). This led to the discovery of first-in-class, drug-like ligands for 9 of 20 targets. This result demonstrates the broad ligandability of WDRs. This extensive resource of reagents and knowledge will enable further discovery of chemical tools and potential therapeutics for this important class of proteins.

## Introduction

WD40 Repeat (WDR) proteins comprise one of the largest protein families with ∼349 WDR encoding genes in the human genome. WDRs are generally understudied (**Fig. 1a**, **Table S1**) though many have strong genetic links to human diseases and are involved in a wide variety of cellular processes including epigenetics, ubiquitin signaling, DNA repair, RNA splicing, and immune system signaling^1^. This family harbors more ‘essential genes’ in cancer than any other protein family (**Fig. 1b, Table S1**)^2^, and is also involved in many neurological and inflammatory diseases. Despite the strong links to disease, and unlike other large target classes such as GPCRs or kinases, the WDR family is largely unexplored with respect to drug discovery.

**Fig 1.**
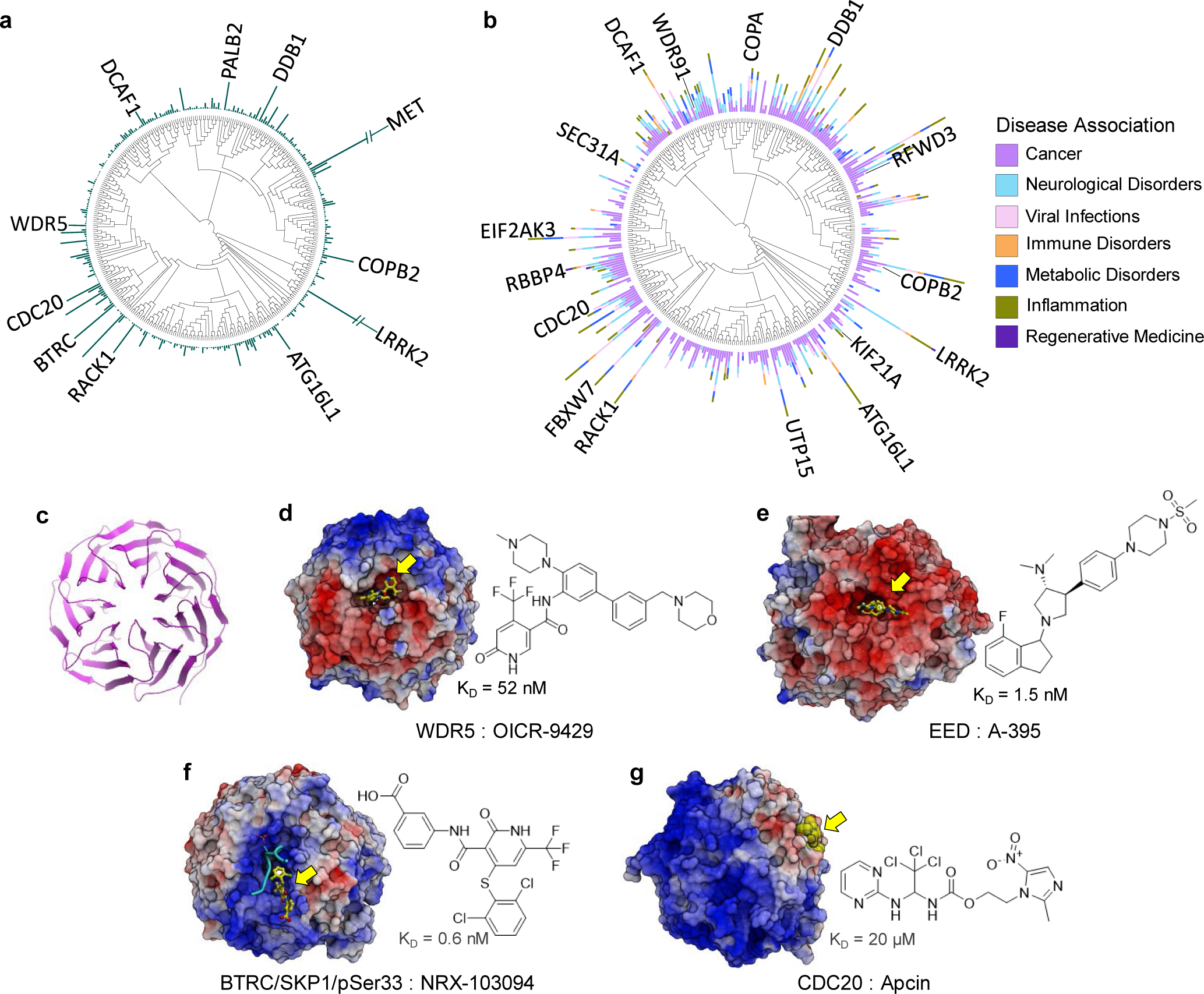
WDR research activity as seen through the lens of ligand-bound structures. A phylogenetic tree annotated with **(a)** the number of publications (reporting on ≤ 30 genes) indexed in PubMed, **(b)** Log_2_ of the number of publications indexed in PubMed associating the protein with the specific disease category, **(c)** A ribbon representation of a typical 7-bladed propeller WDR domain, **(d)** WDR5 in complex with OICR-9429 (PDB: 4QL1)^6^, **(e)** EED in complex with A-395 (PDB: 5K0M)^10^, **(f)** BTRC/SKP1/pSer33 in complex with NRX-103094 (PDB: 6M91)^12^, **(g)** CDC20 in complex with apcin (PDB: 4N14)^13^. These structures illustrate the diversity of electrostatic potential distribution around the protein surface and in the central pocket. Small molecule ligands are displayed in yellow in the 3D structure, and to the right are the chemical structures with the associated binding (K_D_) constants. Surface electrostatic potential with colors saturating below −5 kcal/e.u. charge units (red) and above +5 kcal/e.u. charge units (blue).

The WDR domain is a protein-protein interaction (PPI) module often acting as a substrate recognition or scaffolding subunit within larger multiprotein complexes^1,3,4^. WDR domains all have a canonical donut shape comprising 7-9 β-propellers, each of which contains the namesake 40-residue WD (tryptophan-aspartate) repeat sequence (**Fig. 1c**). Although WDRs have been shown to mediate PPIs through many surfaces of their conserved fold, the central pocket is frequently involved in key functional interactions with peptide regions of partner proteins such as degrons or recognition of post-translational modifications. Furthermore, some WDRs also bind nucleic acids^4,5^.

Interestingly, neither the outer surface of the WDR donut, nor the residues lining the central pocket are conserved across the WDR family (**Fig. 1c-g**). With the exception of a few very closely-related proteins, WDRs are only conserved in their protein fold, but not in their functional interaction sites. Nevertheless, the central pocket often has the appropriate size and physicochemical properties for potent binding to drug-like small molecules. We were among the first to demonstrate that it is possible to disrupt PPIs of WDR proteins with drug-like small molecules that compete with or enhance binding of native interacting proteins/peptides (**Fig. 2**). For example, antagonists of WDR5 can disrupt its interaction with Mixed Lineage Leukemia (MLL) histone methyltransferase (**Fig. 1d**), thereby abrogating MLL function to suppress growth in leukemia and breast cancer^6-9^. Similarly, small molecule antagonism of the central pocket of EED (**Fig. 1e**), a WDR subunit of the polycomb repression complex 2 (PRC2), disrupts binding of PRC2-stimulatory proteins with concomitant inhibition of PRC2 catalytic activity^10,11^.

**Fig. 2.**
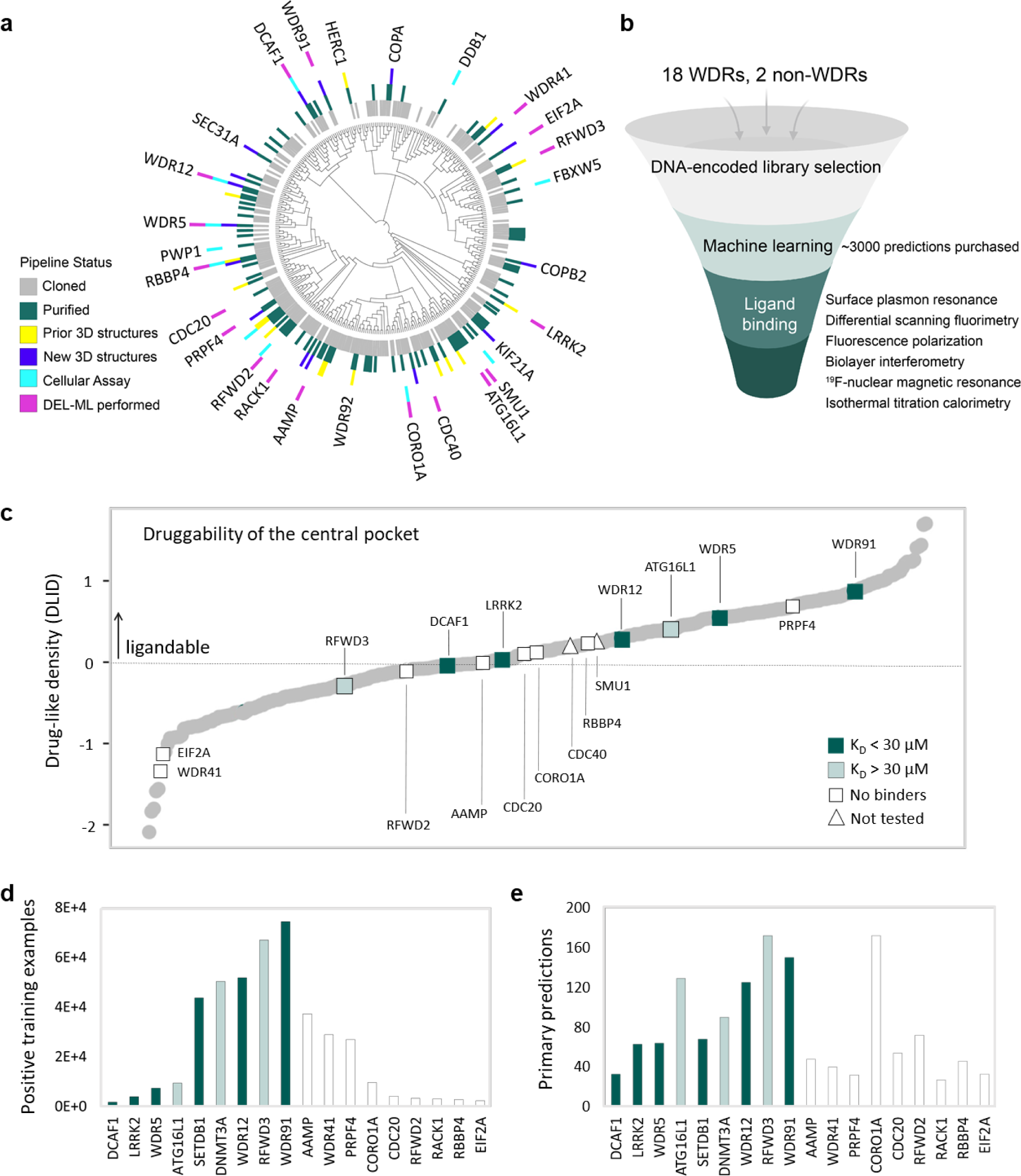
A toolbox for developing pharmacological tools for WDR-containing proteins. **(a)** Phylogenetic tree of the WDR protein family illustrating the breadth of purified proteins, crystal structures, experimental protocols and hit-finding data that this study has made available, (**b**) The hit-finding funnel encompassing DEL selections to ligand binding assays, (**c**) Ligandability of the central pocket in AlphaFold-generated structures^37^ evaluated with the drug-like density index (DLID)^34^ calculated with ICM (Molsoft, San Diego) across all human WDR containing proteins (pockets predicted to be ligandable have a DLID > 0; details in **Table S7**), (**d**) The number of positive training examples calculated from the DEL selections (**Tables S8, S9**), (**e**) The number of primary predicted ligands tested; the complete list (including follow-up predictions) is in **Fig. S1, Table S10**. (Compounds predicted to bind CDC40 and SMU1 were not purchased but are listed in **Table S10**.)

WDR ligands can also act as activators, for example by ‘locking in’ a native binding partner via a molecular glue mechanism such as for BTRC E3 ligase and β-Catenin^12^ (**Fig. 1f**). Small molecule ligands have also been reported to bind (in pockets) on the outer surface of some WDRs, such as the CDC20 E3 ligase (**Fig. 1g**), but such ligands are of lower potency^13^. There is growing interest in ligand discovery for WDRs from the context of proximity pharmacology. For example, the WDRs DCAF1^14-17^, DCAF11^18^, and DCAF16^19^ are substrate recognition subunits of Cullin E3 ubiquitin ligases^20^, and several interact with DDB1^21^.

Notwithstanding these exciting results over the past decade or so, the vast majority of WDR proteins are both understudied in terms of their biological function (**Fig. 1a**), and unexplored with respect to small molecule antagonism. We hypothesized that the WDR protein family harbours many more druggable members, and set out to test this in the context of Target 2035, a global initiative to develop pharmacological tools for all druggable human proteins^22-24^. We were particularly interested in developing scalable hit-finding approaches that could be applied broadly. To this end, we set out to explore the broader applicability of DEL-ML which overcomes many of the practical liabilities of DEL screening^25-27^. This approach was first demonstrated by McCloskey et al.^25^ for a member of each of the highly druggable protein kinase, hydrolase, and nuclear receptor target classes.

In this resource we describe a set of unencumbered reagents and knowledge to enable chemical biology and drug discovery for a wider variety of WDR proteins distributed across the large family. We first expressed and purified ∼95 affinity-tagged recombinant WDR domains, and established a suite of assays to characterize ligand binding and cellular target engagement^28,29^. We then applied DEL-ML hit-finding on a subset of these proteins, discovering seven novel, unique hits for six targets and weak binders for an additional three targets. A high-throughput crystallography workflow^30^ yielded 21 apo and ligand-bound 3D WDR structures with which to interpret protein-ligand interactions. The results indicate that many understudied WDR proteins are indeed ligandable with drug-like small molecules, and that DEL-ML has broader applicability across the diverse, non-conserved WDR binding pockets. These data, methods and the related reagents are a significant resource to the community that will enable the development of pharmacological tools for WDR proteins to further explore their biology, interaction networks, and therapeutic potential.

## Results

### WDR domain protein production and 3D structures

To choose a test set of WDR proteins, we considered proteins that were both strongly linked to diseases such as cancer (**Fig. 1**), as well as those that were not well characterized, including many for which recombinant protein or experimentally determined 3D structures have not been reported (**Table S1**). A total of 259 WDR proteins – more than half the 349-member WDR family – were selected for recombinant expression and purification (**Tables S2, S3**).

Previous experience in our group suggested that recombinant expression of human WDR domains in a Baculovirus Expression Vector System (BEVS) was often more successful than expression in *E. coli*. We therefore designed on average two expression constructs for each of the WDRs and subcloned these for baculovirus-mediated expression in Sf9 cells. Because of conservation of the overall WDR fold, we were able to readily predict the domain boundaries, including the frequent addition of helices or small loops and folded regions internally or at the immediate termini, using homology modeling; this initial phase of the project predated the release of AlphaFold^31-33^. Of the 259 proteins tested for expression, 95 were purified at scale with at least one affinity tag, and 25 with a luminescent tag (**Fig. 2a, Tables S2, S3**).

Quality control of the recombinant protein included using SDS-PAGE and mass spectrometry to confirm purity (> 90%) and the expected molecular weight, respectively. Finally, differential scanning fluorimetry (DSF) (**Table S4**) was performed to ensure each purified protein was stable and folded in the absence of its interaction partner(s). These data showed that a large proportion of WDR domains can be produced in isolation for further study – including many that are central components of large macromolecular complexes. Protein crystallization was attempted for select WDRs that had sufficient yields to setup at least one 96-well crystallization tray and homogeneous elution profiles on size exclusion chromatography. Overall, 14 apo WDR structures were generated to support structure-guided drug discovery efforts (**Fig. 2a, Tables S5, S6**). The protocols for expressing, purifying and crystallizing the apo WDR domains greatly facilitated the rapid characterization and advancement of ligands identified through DEL-ML as described below^14,27^.

Upon release of the AlphaFold structure database^32^, we analyzed the electrostatics and central pocket ligandability of all 349 human WDR-containing proteins with predicted structures. The use of AlphaFold predictions to map electrostatics and ligandable pockets in WDRs was supported by our observation that less than 1.3 Å backbone RMSD separates the predicted WDR structures of UTP15 or EIF2A and their subsequently released crystal structures (PDB IDs: 7RUO, 8DYS). Overall, about half of the central pockets were predicted to be ligandable^34^ (drug-like density (DLID) > 0, **Fig. 2c, Table S7**), which we believe is a conservative estimate considering that pockets are sometimes occluded in the absence of ligands. For example, the central pocket of the apo structure of EED is occluded by several residues which are displaced by binding of drug-like small molecule ligands^10,35^. The diversity in shape and electrostatics of the central pocket was striking^36,37^. In fact, it was previously shown that structural diversity is even observed between two WDR binding sites targeted by the same endogenous ligand. For example, an arginine occupies the structurally dissimilar central pockets of the epigenetic regulators WDR5 and RBBP4^3^ yet WDR5 ligands do not bind to RBBP4. This analysis suggests that the discovery of selective ligands for a given WDR protein of interest is quite feasible.

### High throughput DEL selection reveals significant ligandability of WDRs

DEL selections were performed for 18 N-terminally biotin- or his-tagged WDRs (**Fig. 2b, Table S8**), 17 of which had no reported small molecule ligands at the start of this program. WDR5 was included in the screen as a positive control, as it is druggable and has elicited clinical interest in the context of treating leukemias^6,8,35^. Given the novelty of the WDR family as a target class, we also included the PWWP domain of DNMT3A and the triple Tudor domain (TTD) of SETDB1 as two unrelated PPI domains that have been shown to be ligandable^38-40^.

The DEL deck used for these screens comprised compounds synthesized by applying a split-and-pool methodology^25^. This approach allows multiple thousands of chemical building blocks to be reacted with each other in a combinatorial manner to access multiple billions of diverse and distinct compounds. This diverse set of DNA-encoded compounds encompassed common structural motifs optimized for small molecule drug discovery. Screening of the DEL deck was accomplished by the affinity-mediated selection of a mixture of all available libraries. Captured targets and associated library members were stringently washed under native conditions followed by the recovery of retained library members by thermal denaturation of the target. After two cycles of selection and recovery, library members were amplified by PCR and sequenced to generate a minimum of one million reads per library per condition.

After parallel selection outputs were sequenced, normalized, amplification duplicates removed, sequence identifiers translated back into chemical and condition identifiers, and the statistical significance of all building block combinations calculated in all conditions, the quantity and quality of the enriched compounds was assessed. Featurized structural Information of enriched building block combinations resulting from each of these selections formed the input for test and training sets for machine learning algorithms. These inputs are referred to as Positive Training Examples (PTEs) (**Fig. 2d, Table S9**). Models that performed well on the test set were then used to predict the binding of commercially available compounds from large virtual libraries such as Enamine REAL and Mcule databases^14,25,27^. Depending on the number of predictions and in-stock availability, between 33 and 200 predicted compounds were purchased for 18 targets (16 WDRs) totaling ∼3000 compounds in all (**Figs 2e, S1, Table S10**). Due to logistical reasons, the predictions for two targets (CDC40 and SMU1) were not purchased or tested but are listed in **Table S10**.

Each compound was evaluated for solubility and aggregation up to (at least) 200 μM by dynamic light scattering (DLS)^41-43^, to identify potential sources of assay interference. Soluble compounds were assayed for binding to their predicted target using a variety of biophysical and biochemical methods. For this, we chose surface plasmon resonance (SPR) as the primary assay for hit validation as it is a generic, well-developed method for detecting small-molecule interactions with a protein immobilized on a biosensor chip via an affinity tag. SPR has good throughput and can be run in a screening mode for many compounds or in a dose-response mode to determine the affinity of individual compounds (**Table S10**).

We found that most WDRs are readily assayed by SPR under a limited set of standard buffer conditions (see Methods). Compounds that appeared to bind to their predicted target by SPR were then assessed in dose response mode. Compounds with apparent affinities below ∼200 μM K_D_ value were advanced to an orthogonal assay, such as DSF, Isothermal titration calorimetry (ITC), ligand-observed ^19^F-NMR, or biolayer interferometry (BLI), depending on the target and ligand (**Fig. 2b**). We discovered seven hits (K_D_ < 30 μM) for five WDRs and for SETDB1 (**Figs. 3-8**). We also discovered a weak, but detectable binder (K_D_’s ranging from 40-200 μM) for each of three additional targets, ATG16L1, RFWD3, and DNMT3A (**Table S10**). We subsequently employed X-ray crystallography to characterize the binding sites of protein-ligand complexes. In some cases, the binding site of the ligand could also be identified by a peptide displacement assay using fluorescence polarization (FP). Importantly, the identified ligands were all novel chemotypes for their respective targets.

**Fig. 3.**
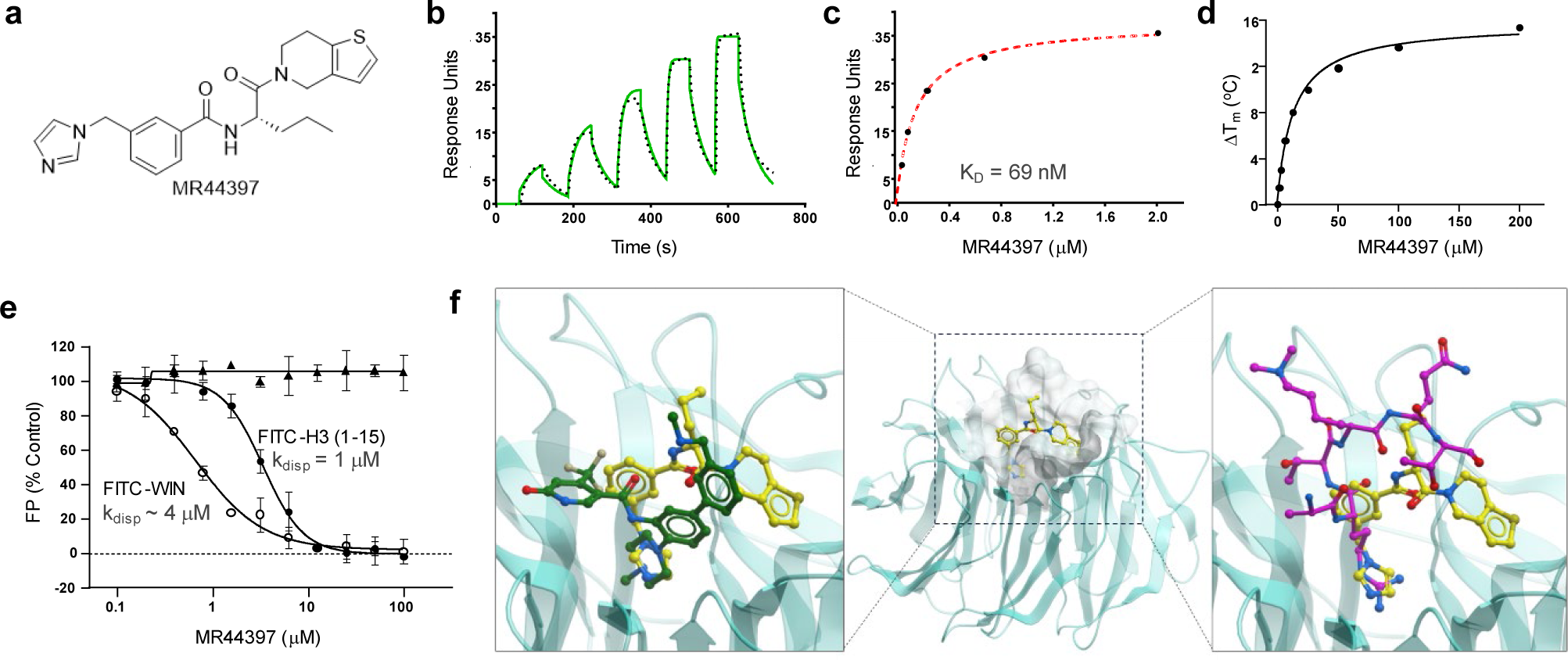
Characterization of a novel WDR5 ligand. **(a)** Chemical structure of MR44397. **(b)** An SPR sensorgram representing the kinetic fit (black dots), K_D_ = 69 nM, **(c)** SPR steady-state response (black circles) using a 1:1 binding model. **(d)** Concentration-dependent protein thermal stabilization determined for MR44397; ΔT_m_, calculated from fitting transition curves using the Boltzman sigmoid function, is plotted against concentration (see also **Fig. S2a**). **(e)** Fluorescence polarization-based displacement assays shows that MR44397 displaces the FITC-H3 (1-15) (○) with k_disp_ = 1 µM, and the FITC-WIN peptide (● GSARAEVHLRKS) with k_disp_ = 4 µM, but not FITC-RBBP5 peptide (▴, EDEEVDVTSV) which binds at a different site. **(f)** (**Center**) Co-crystal structure of WDR5 (cyan) in complex with MR44397 (yellow sticks, PDB ID: 8T5I); (**left**) a close up view of the MR44397 binding site showing superimposed structures of WDR5 (cyan) in complex with MR44397 (yellow sticks) and WDR5 in complex with the potent WDR5 antagonist OICR-9429 (green sticks, PDB ID: 4QL1)^6^; and (**right**) superimposed structures of WDR5 (cyan) in complex with MR44397 (yellow sticks) and WDR5 in complex with a histone H3K4 peptide (magenta sticks, PDB ID: 2O9K)^44^.

### DEL-ML delivers chemotypes that are chemically tractable

The predicted ligands are chosen from the Enamine REAL and MCule databases. As such, all hits comprise chemical templates that are readily amenable to structure-activity exploration and structural optimization (**Tables 1, S10**). For example, a novel chemotype was discovered for the highly druggable WDR5^35^. Of 64 ML-predicted compounds, nine had SPR K_D_ values less than 100 μM with some SAR (**Table S10**). Two of the most potent compounds are racemates featuring an imidazole and/or tetrahydrobenzo-thiophene or pyridine. The most potent ligand MR43378 (K_D_ = 16 μM) was subjected to chiral separation and subsequent testing by SPR showed that the *S-*enantiomer (MR44397) had a K_D_ of 69 nM (**Figs. 3a-c**).

A suite of orthogonal assays, including DSF, FP, and BLI, confirmed potent binding (**Figs. 3, S2**). MR44397 substantially stabilized WDR5 as measured by DSF (**Figs. 3d, S2a**), and displaced two central pocket-binding peptides (histone H3K4 and WIN peptides)^44^ with K_disp_ of 4 µM, and 1 µM, respectively (**Fig. 3e**). Importantly, MR44397 did not displace an RBBP5 peptide (aa 371-380), which binds on a surface outside the central pocket of WDR5 (**Fig. 3e**)^45^. To gain structural insights into the binding mode of the compound, we determined the co-crystal structure of MR44397 in complex with WDR5, referred to here as WDR5-MR44397 (**Tables S5, S6**). The compound occupies the central pocket of WDR5 (**Fig. 3f**), in agreement with the FP displacement data (**Fig. 3e**). These data clearly demonstrate a new chemically tractable series for WDR5, which modulates functional protein-interactions of the target.

For DCAF1, only a small set of 33 commercial compounds were initially screened. As recently reported, the most potent compound Z1391232269 (compound 3c)^14^, had K_D_ of 14 µM and binding was confirmed by multiple orthogonal methods (**Figs. 4, S3**). Further investigation and structural analysis showed that the *S-*enantiomer (compound 3d/OICR-6766)^14^, which was present as a 4% impurity in the commercial material, bound to DCAF1 with a K_D_ of 490 nM, while the *R*-enantiomer had a K_D_ of 14 µM (**Table 1**, **Fig. 4**). OICR-6766 binds deep inside the WDR central pocket (**Fig. 4d**, PDB: 7UFV, 8F8E) with both aromatic rings nestled in hydrophobic pockets.

**Fig. 4.**
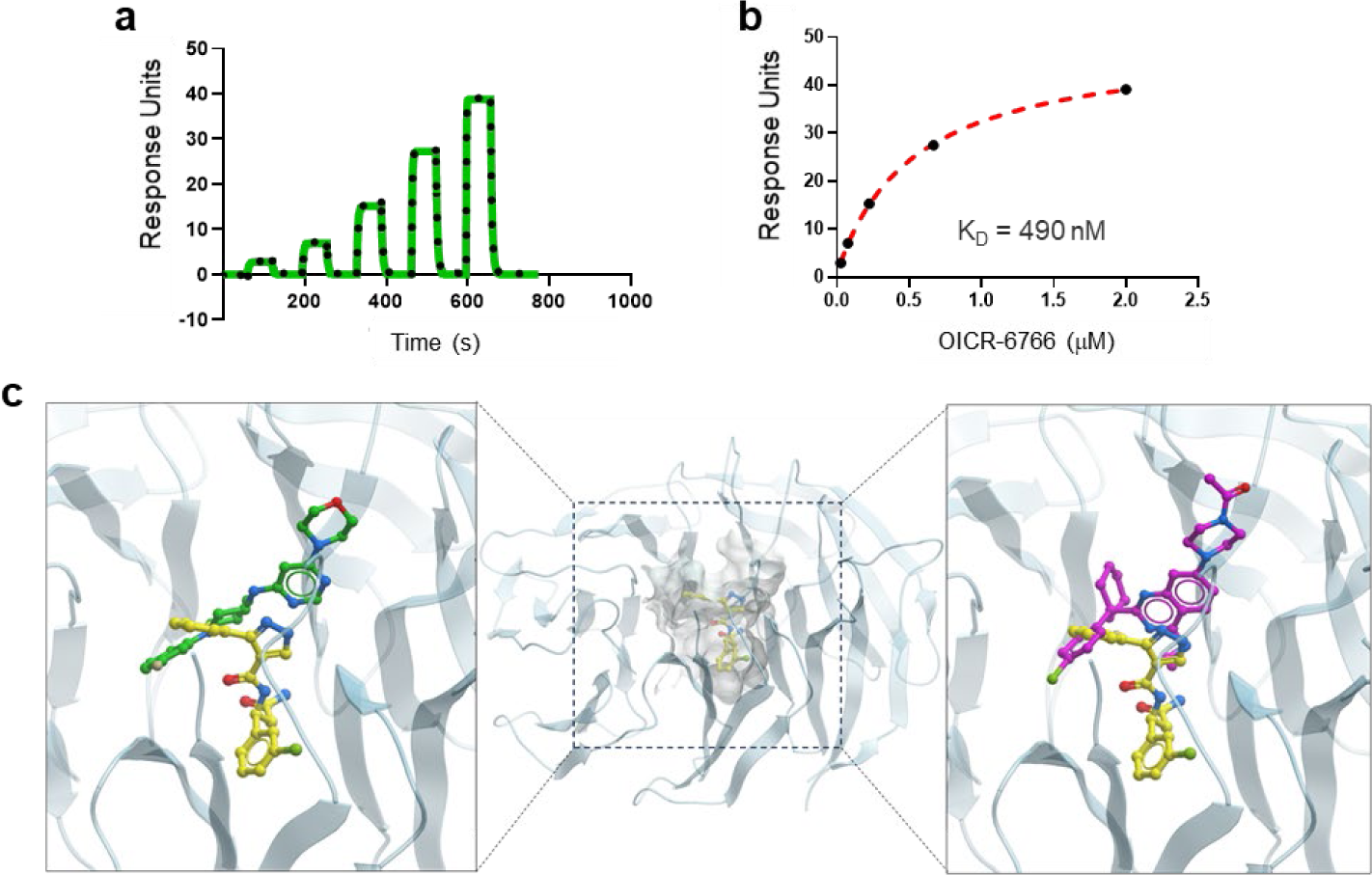
A potent DCAF1 ligand is nestled deep inside the central pocket. **(a)** A representative SPR sensorgram for OICR-6766 (K_D_ = 490 nM) with the kinetic fit (black dots), **(b)** SPR steady state response (black circles) using a 1:1 binding model, **(c)** (**Center**) Co-crystal structure of DCAF1 (cyan) in complex with OICR-6766 (yellow sticks, PDB ID: 7UFV)^14^, (**left**) superimposed structures of DCAF1 (cyan) in complex with OICR-6766 (yellow sticks, PDB ID: 7UFV) and DCAF1 in complex with CYCA-117-70 (green sticks,PDB ID: 7SSE)^46^, and (**right**) superimposed structures of DCAF1 (cyan) in complex with OICR-6766 (yellow sticks,PDB ID: 7UFV), and DCAF1 in complex with compound 13 (magenta sticks, PDB ID: 8OO5)^16^.

**Table 1.**
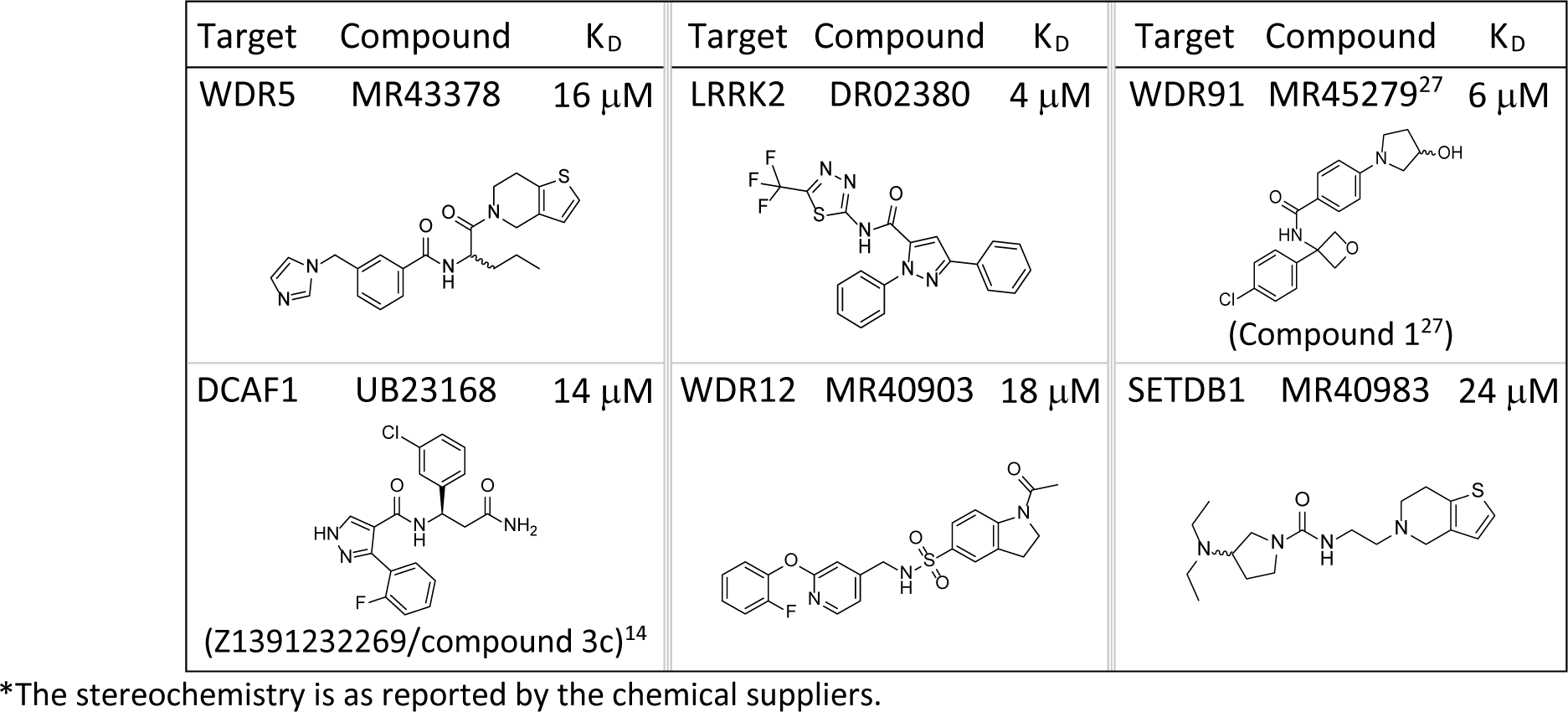
Chemical structures and (SPR) binding affinities for hits. (The optimized hits are presented in Figures 3-9.)

Following the initial set of 33 compounds (**Table S10**), an additional ∼547 compounds were purchased and tested as additional hit identification or hit expansion exercises to the original hits, utilizing a variety of methods including DEL-ML GCNN model scoring, virtual docking, 2D similarity and shape based (ROCS) searching the commercially available EnamineReal space. While these data gave some hints of the SAR around the OICR-6766 chemotype, there were no compounds that were significantly more potent. Instead, structure-guided drug design was used to optimize OICR-6766 into OICR-8268 (K_D_ = 38 nM) that engages DCAF1 in cells^14^, demonstrating the chemical tractability of the initial hit from only 33 predicted compounds. Notably, inspection of the DEL selection output data confirmed that the building block pair corresponding to the hit exists in the DEL deck, but only as the *S*-enantiomer, and was enriched by DCAF1. Interestingly, contemporaneous with our studies^14^, novel ligands for DCAF1 were discovered using computational screening methods^46^ and HTS^16^. Both reported ligands bind closer to the outer surface of the central pocket, and taken together with these results indicate that DCAF1 is a fairly druggable WDR.

First-in-class ligands for the WDR domain of LRRK2, and WDR12 were also discovered from DEL-ML predictions. Of ∼70 predicted compounds tested for LRRK2, two hits with unique chemotypes were identified (**Table 1**, **Fig. 5**). Both compounds bind in a dose-dependent manner, but the level of binding is notably less than ideal. DR02380 binds to LRRK2 with an estimated K_D_ value of 4 µM (**Fig. 5**, **Tables 1, S10**). Taking advantage of the trifluoromethyl group, we used ^19^F-NMR as an orthogonal method to confirm binding (**Fig. 5c**). The second chemotype, DR02034 (**Fig. 5d-f**), has an estimated K_D_ value of 11 µM. Its binding was also confirmed by ^19^F NMR of a fluorophenyl analog DR02244 (**Fig. 5d, g-i**).

**Fig. 5.**
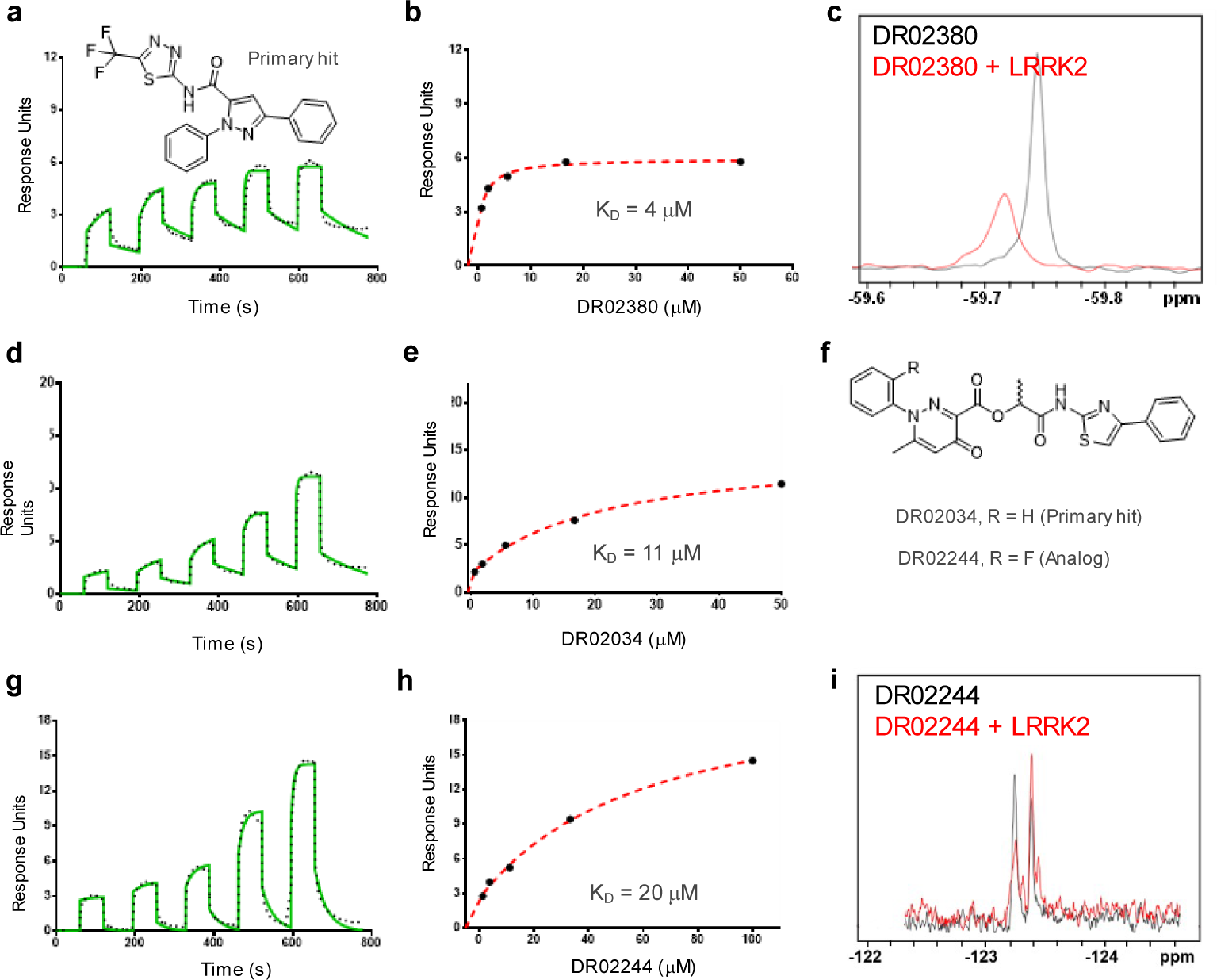
Two unique chemotypes were discovered for the WDR domain of LRRK2. **(a,b)** SPR sensorgram with the kinetic fit and steady state response for binding to DR02380, **(c)** (ligand observed) ^19^F-NMR spectrum of 10 μM DR02380 in the absence (black) and presence (red) of 20 μM LRRK2 WDR domain, demonstrating a significant change (in peak height and broadening) in the chemical environment of the trifluoromethyl group, **(d,e)** SPR sensorgram with the kinetic fit (black dots) and steady state response (black circles) for binding to DR02034, **(f)** Chemical structures of DR02034 and a fluorine analog DR02244, **(g,h)** SPR sensorgram with kinetic fit and steady state response for binding to DR02244, **(i)** (ligand observed) ^19^F-NMR spectrum of 10 μM DR02244 in the absence (black) and presence (red) of 20 μM LRRK2, demonstrating a small but noticeable change (in intensity and splitting) in the chemical environment of the fluorine atom. (A 1:1 binding model is used in b, f, h.)

For WDR12, we tested 530 predicted ligands, for which 39 showed dose-response binding (with K_D_ < 100 μM) by SPR. Interestingly, the majority of these ‘hits’ shared structural features including methyl sulfonamides or methyl sulfones, bi-aryl or aryl cyclohexyl ethers, providing limited SAR (**Table S10**). MR40903 is the highest affinity ligand (K_D_ = 18 μM) which harbors a fluorine atom (**Table 1**) which, when replaced with chlorine yielded MR44915 with slightly better affinity (K_D_ = 4 μM). (**Fig. 6a-c**). Dose-response binding of MR40903 was confirmed using ^19^F-NMR (**Fig. S4**), while hydrogen-deuterium exchange mass spectrometry (HDX-MS)^47^ confirmed binding to the more potent MR44915 (**Fig. S5**). HDX-MS of WDR12 in the presence of MR44915 revealed regions around the central pocket (shaded in blue) that were protected from deuterium uptake. Moreover, these protected regions included three peptides on the ‘top’ of the WDR fold that point toward the central pocket (**Figs. 6d**, **S5**). These data suggest that MR44915 likely binds near the top of the ‘donut’.

**Fig. 6.**
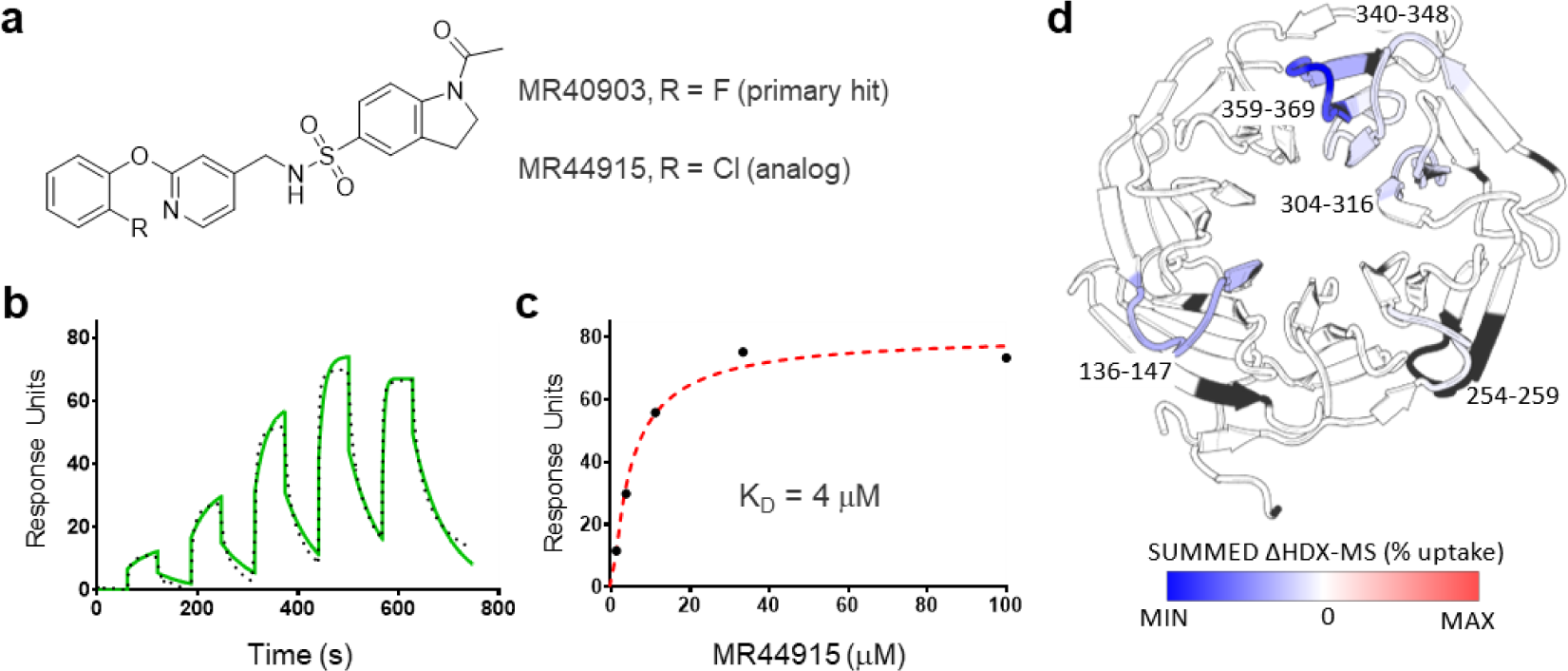
A novel ligand for WDR12 appears to bind near the central pocket. **(a)** Chemical structure of MR40903 (DEL-ML hit) and its analog MR44915 (**Tables 1**, **S8**), **(b,c)** A representative SPR sensorgram with the kinetic fit (black dots) and SPR steady state response (black circles) using a 1:1 binding model, **(d)** A cartoon representation of apo WDR12 (PDB: 6N31) with shading representing deuterium uptake in the HDX-MS experiment. The regions shaded in blue have the least rate of deuterium uptake. There was no sequence coverage for the regions shaded in black (with additional details in **Fig. S5**).

DEL-ML screening of WDR91 yielded 150 ML-predicted compounds from which we discovered a first in class, potent ligand MR45279^27^ (K_D_ = 6 µM) (**Fig. 7a-c, Table S10**), which readily co-crystallized (PDB: 8SHJ). Contrary to the hits for DCAF1^14^, WDR5 and possibly WDR12, MR45279 binds in a side pocket nestled between two beta propeller blades and located at the bottom side of the WDR domain (**Fig. 7c**). Importantly, the WDR91-MR45279 structure demonstrates that side pockets, which are often additional functional interaction sites in WDRs, can also be targeted by small-molecules with good (6 µM) affinity.

**Fig. 7.**
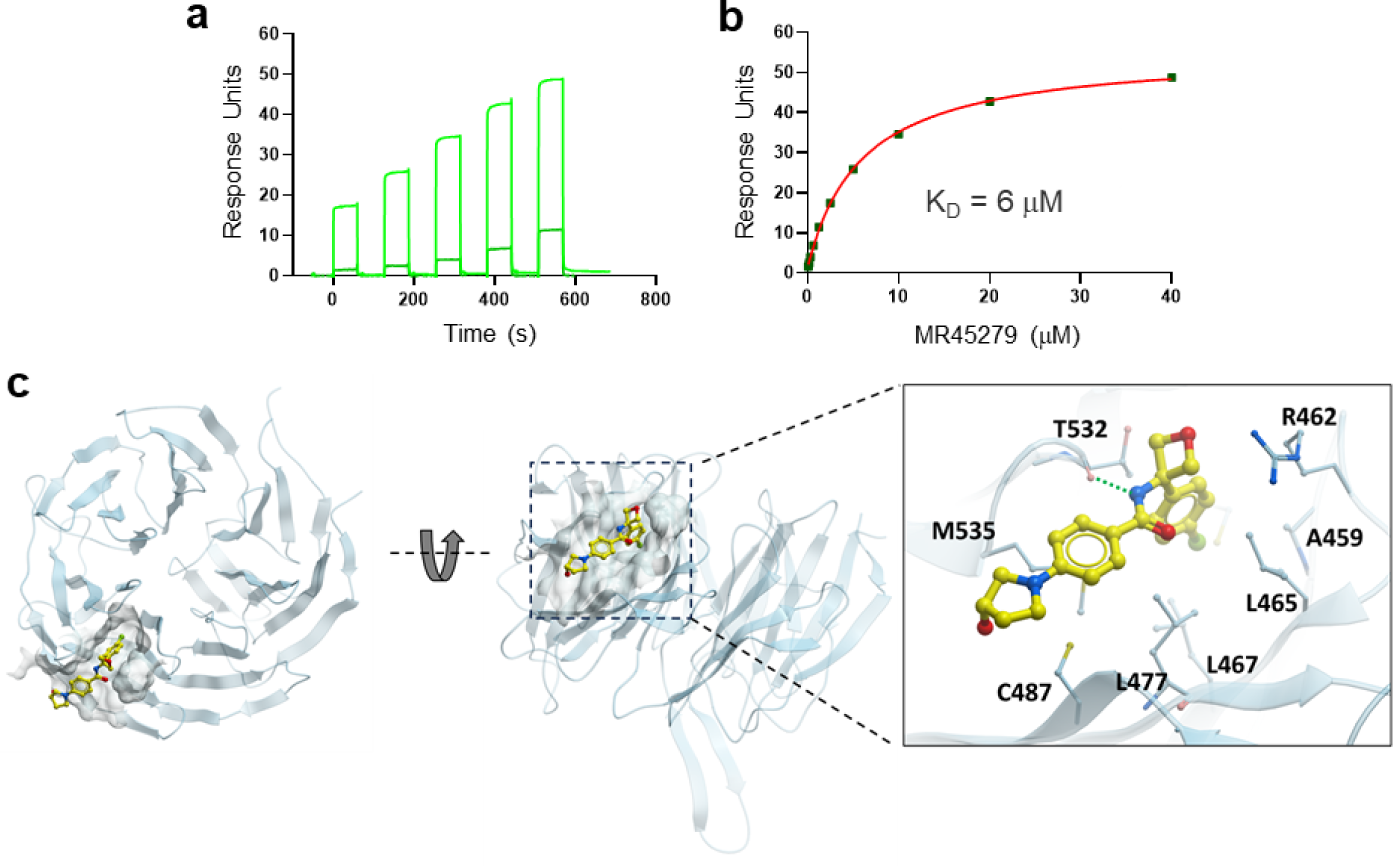
WDR91 first-in-class ligand binds in a side pocket. **(a)** representative SPR sensorgram for MR45279 (Table 1, compound 1^27^), **(b)** SPR steady state response (black circles) using a 1:1 binding model, **(c)** (**Left**) Cartoon representation of WDR91 bound to MR45279 (yellow sticks) in the side pocket of the WDR domain, and (**right**) zoomed in view highlighting the key residues interacting with MR45279. The hydrogen-bonding interaction between the backbone oxygen of threonine (T532) and the amide nitrogen of the compound is shown with a dotted green line.

Finally, for SETDB1 TTD, which recognizes lysine methylation and acetylation marks on histone H3^48^, and for which ligands have previously been reported^39,40^, testing of 68 ML-predicted compounds using an FP-based peptide displacement assay yielded novel hits (**Table S10**).Orthogonal confirmation of binding using SPR identified MR40983 (a racemate) as the most potent ligand with K_D_ value of 24 μM (**Tables 1, S10**). MR40983 is a medium MW compound (350 Da) containing two tertiary amine moieties.

Using these preliminary data, another ∼107 compounds were procured using SAR-by-catalog (**Table S10**) and similarly tested in a peptide displacement assay and then by SPR. Three hits - MR43625, MR43615, MR43579 – were identified with the most potent MR43625 having a K_D_ of 12 μM (**Table S10**). All three compounds were higher MW (400-450 Da) and contained two tertiary amines (mimics of methylated lysines). The most potent hit MR43625 (a racemate) was co-crystallized with SETDB1-TTD (PDB ID: 8UWP). The structure revealed that MR43625 binds at the interface between the TD2 and TD3 by inserting one tertiary amine moiety into each of the aromatic cages of TD2 and TD3, and that there is a better fit to the *S-*enantiomer (**Fig. 8e**). To confirm this observation, the enantiomers were procured and tested by SPR. The *S-*enantiomer, MR46747, binds with a K_D_ value of 4 μM while the *R-*enantiomer binds with a K_D_ value of 70 μM (**Figs. 8b, c, Table S10**). Further, MR46747 displaces H3K9me2K14Ac with k_disp_ of 36 μM.

**Fig. 8.**
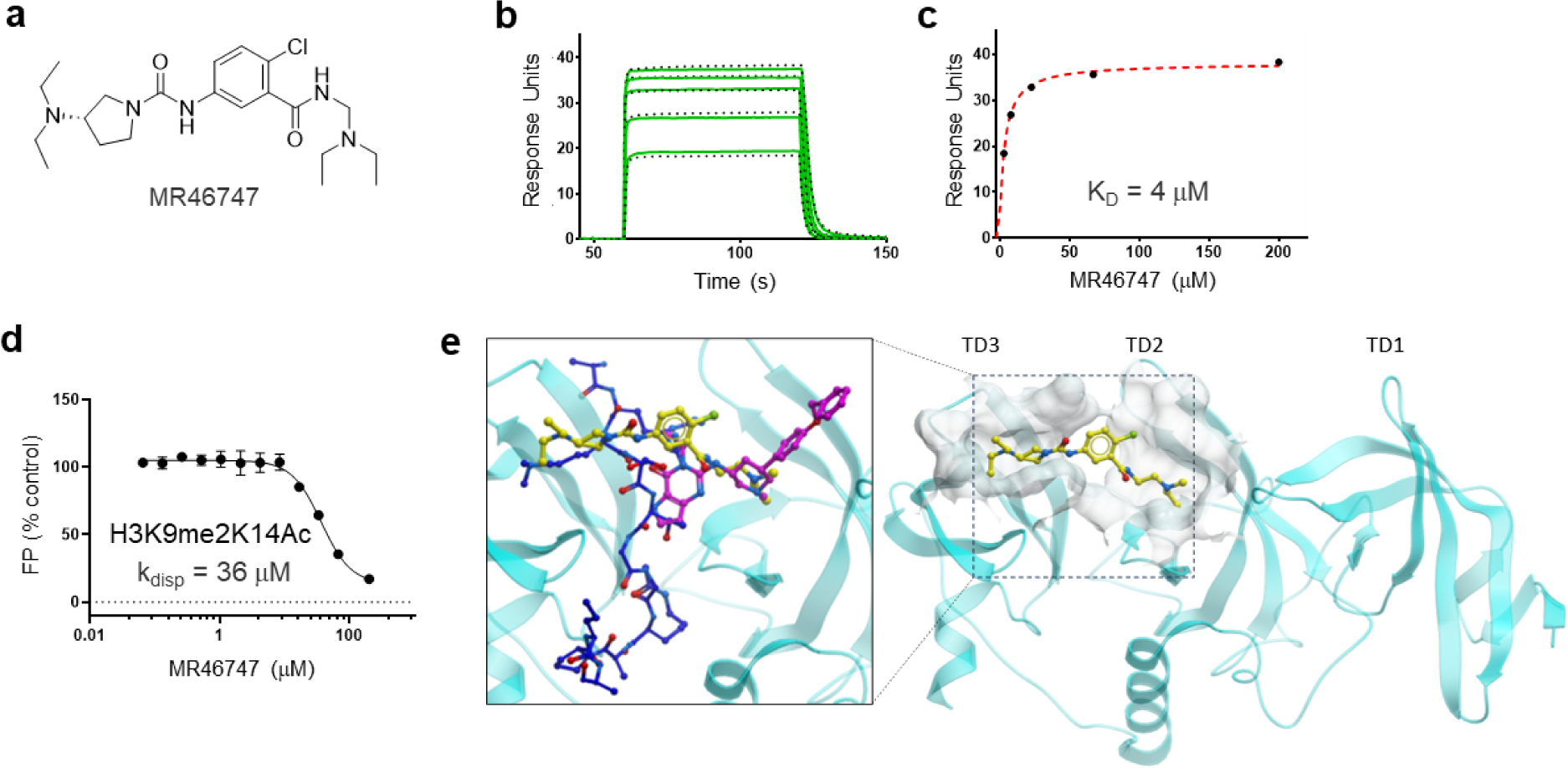
A novel peptide-competitive ligand of SETDB1-TTD. **(a)** Chemical structure of MR46747 (the S enantiomer of the DEL-ML hit MR43625), **(b)** A representative SPR sensorgram with the kinetic fit (black dots), **(c)** SPR steady state response (black circles) using a 1:1 binding model, **(d)** FP-based displacement assays shows that MR46747 displaces the FITC-H3K9me2K14ac (1-25) (●) with K_disp_ = 36 µM, **(e)** (**Right**) Co-crystal structure of SETDB1-TTD (cyan) in complex with MR46747 (yellow sticks, PDB ID: 8UWP), and (**left**) superimposed crystal structures of SETDB1-TTD (cyan) in complex with MR46747 (yellow sticks, PDB ID: 8UWP), SETDB1-TTD in complex with H3K9me2K14Ac (blue, PDB ID: 6BHD), and SETDB1-TTD in complex with (*R*, *R*)-59 (magenta, PDB ID: 7CJT).

### Newly discovered hits are selective

To assess the selectivity of the identified hits, we used SPR to evaluate binding to six targets. For each compound the percent binding of each ligand to each of the six proteins was measured in triplicate, at a compound concentration of 50 μM. All hits exhibited selectivity for their intended targets (**Fig. 9, Table S11**) with over 77% binding observed except for the LRRK2 hits, which displayed lower binding levels to LRRK2 (10-20%) and showed no binding to the remaining five targets. Lower percent binding to LRRK2 may be due to the apparent poorer solubility of the LRRK2 compounds (**Table S10)**.

**Fig. 9.**
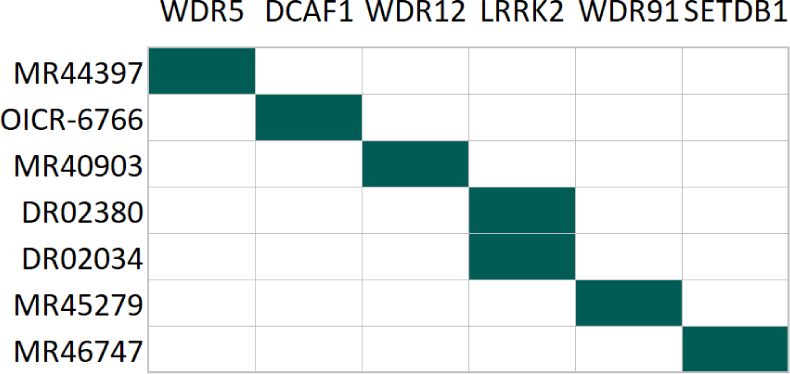
Selectivity profile of optimized hits discovered by DEL-ML. For each ligand (in rows), the green color indicates the target (in columns) with highest percentage binding relative to other targets. The experiments were performed in triplicate at a compound concentration of 50 μM (**Table S11**).

### Cellular assays to test compound activity

An important step in the drug discovery process is confirming target engagement of a small molecule in a native cellular environment. To develop specific cellular assays for individual WDR proteins we utilized the NanoLuc Luciferase (NL) technology, including HiBiT Cellular Thermal Shift Assay (HiBiT CETSA), NanoBRET-based tracer assays, as well as protein-protein interaction (PPI) NanoBRET assays^49^. Below we describe the suite of cell assays developed for WDRs and provide examples demonstrating target engagement by WDR5 ligands (**Fig. 10**).

**Fig. 10.**
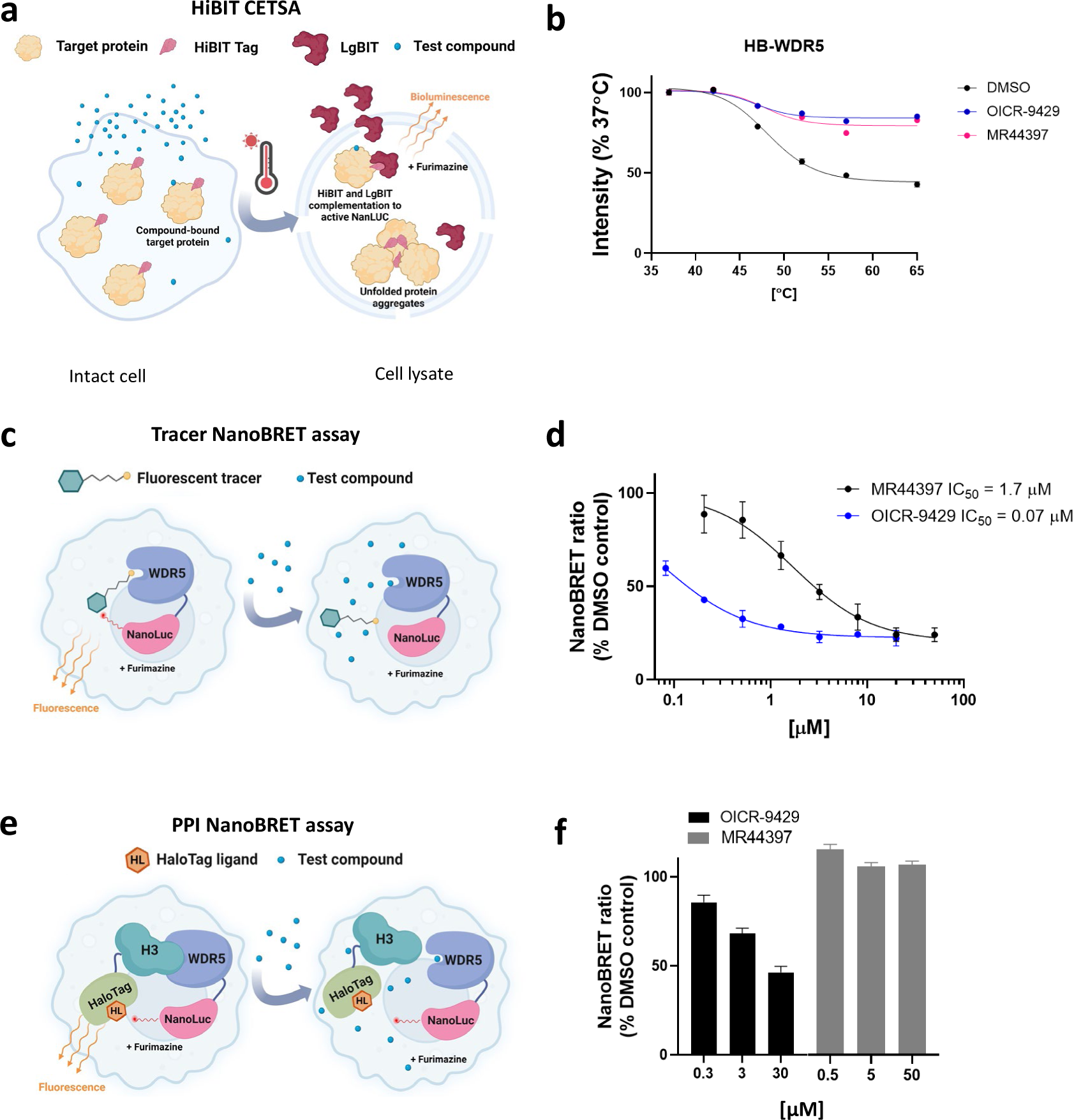
MR44397 cellular WDR5 target engagement. **(a)** Schematic representation of CETSA^28^. **(b)** MR44397 and the WDR5 chemical probe OICR-9429 (positive control) stabilize WDR5 in HEK293T cells. Cells were transfected with N-terminally HiBiT (HB) tagged WDR5 and incubated with 50 µM of MR44397 and 20 µM of OICR-9429 for 1 h. After heating for 3 min in an indicated temperature gradient, cells were lysed and incubated with LgBIT. The results are MEAN+/- SD of NanoLuc bioluminescence signal (n=4). **(c)** Schematic representation of fluorescent tracer nanoBRET assay^29^**. (d)** Dose response of OICR-9429 and MR44397 competition with a fluorescent tracer compound indicated by decreased nanoBRET ratio of WDR5 and tracer interaction^54^ in HEK293T cells. Cells were transfected with N-terminally NL-tagged WDR5 for 24 h and incubated with 1 µM fluorescent tracer and indicated compounds concentrations for 2 h MEAN+/- SD (n=4). **(e)** Schematic representation of a nanoBRET-based protein-protein interaction assay. **(f)** In contrast to the WDR5 chemical probe OICR-9429 (positive control), MR44397 does not decrease the nanoBRET ratio between WDR5 and histone H3 in HEK293T cells. Cells were co-transfected with C-terminally NanoLuc (NL) tagged WDR5 and C-terminally HaloTag (HT) tagged histone H3 for 24 h and incubated with indicated compounds concentrations for 4 h. The results are MEAN+/- SD (n=4).

To measure the effect of small molecule binding on protein thermal stability, we used a split NanoLuc reporter system (HiBiT CETSA)^50,51^. In the assay, the target protein, which is fused to the 11 amino acid HiBiT tag, is overexpressed in cells which are then subjected to a thermal challenge followed by complementation with LgBIT comprising the remainder of the NanoLuc enzyme thus allowing the level of soluble protein to be quantified from the resulting luminescence signal (**Fig. 10a**). As shown in **Figure 10b**, the newly discovered ligand MR44397 and the chemical probe OICR-9429 thermally stabilized exogenously expressed HiBiT-WDR5 in HEK293T cells indicating cellular permeability and binding to WDR5.

To assess the suitability of this assay for the wider collection of WDR proteins we first compared our systematic DSF data on WDR protein melting temperatures (T_m_) (**Table S4**) and the T_m_s for several HiBiT-tagged WDR proteins transfected in HEK293T cells (**Table S12**) relative to those predicted in the Meltome database^52^. This comparison indicates a reasonable agreement between endogenous protein T_m_ and transfected HiBit-tagged WDR protein T_m_. For example, RBBP5 has a high (∼60°C) T_m_ in the Meltome, HiBit CETSA, and DSF (**Tables S4, S12**). Since temperatures above 60-65 °C can impact cellular permeability, the method is not suitable for proteins with high melting temperatures. RBBP4, RBBP5, WDR41, or WDR82 are examples of such WDR proteins (**Tables S4, S12**). In other cases, compound binding did not result in protein stabilization of the full-length protein making the HiBiT CETSA assay not suitable for cellular target engagement. This problem may be resolved by utilization of targeted domains instead of full-length proteins. For example, OICR-8268 potently stabilized the DCAF1-WD40 domain in DSF and HiBiT CETSA assays but not full-length DCAF1^14^.

Another way to measure target engagement in cells is a tracer-based NanoBRET assay using bioluminescence resonance energy transfer (BRET) from NL to a fluorescent tracer molecule^53^. In this assay, the endogenously tagged or exogenously expressed NL-tagged protein of interest acts as an energy donor in the presence of the furimazine substrate, and a cell-permeable small molecule ligand of the target protein covalently conjugated to the fluorescent dye (tracer) acts as an energy acceptor (**Fig. 10c**). The competition of the test compound with the tracer for binding to the protein of interest results in a decrease in NanoBRET signal between the protein and a tracer and allows the determination of cellular permeability, apparent binding affinity, and residence time of the test compound in intact as well as permeabilized cells^53^.

As an example, the IC_50_ determination in the tracer NanoBRET assay is demonstrated using the WDR5 tracer (**Fig. 10d**)^54^. Both MR44397 and the chemical probe decreased the NanoBRET ratio in a concentration-dependent manner with expected different potencies. This assay confirmed cellular permeability and target engagement of MR44397 with an IC_50_ of 1.7 µM.

In cases where the WDR protein has a known interaction partner that is expected to compete with the ligand of interest, cellular activity and target engagement can be tested with a PPI NanoBRET assay^55,56^. In the PPI NanoBRET assay, one of the proteins of interest is tagged with NL (energy donor), and the other protein of interest is tagged with HaloTag (HT) that binds covalently to a fluorescent chloroalkane ligand and acts as the BRET acceptor if the two proteins are in close proximity (**Fig. 10e**). Besides the measurement of the interactions between full-length proteins, the assay can be modified to measure the BRET ratio between short peptides or isolated domains^29^. An advantage of using individual domains is that the NL and HT tags are likely to be closer upon protein interaction with a resultant higher BRET signal compared to larger full-length proteins where the tags may be more distal to the interaction site.

**Figure 10f** shows the PPI NanoBRET assay for the WDR5-Histone H3 interaction competed with MR44397, and the chemical probe OICR-9429. Although both compounds bind to the central pocket of WDR5 and displace H3 peptide in vitro (**Fig. 3**), only OICR-9429, which is 20 times more potent than MR44397 in the tracer NanoBRET assay (**Fig. 10d**) was able to decrease the NanoBRET ratio. This result exemplifies our observations that compounds required to observe a robust disruption of a PPI in general display greater potencies.

We have developed several additional PPI NanoBRET assays for WDRs which will be useful as potent ligands are developed in the future (**Table S13**). If possible, to further validate the assay and to determine the assay window, we also utilized genetic mutations in donor or acceptor proteins known or predicted to disrupt the interaction. For example, a previously published R29D mutation within CORO1A^57^ resulted in a decrease in NanoBRET ratio with ACTB by 50% and a double mutation predicted from a co-crystal structure of FBXW7 (R465E, R505E) completely abolished the NanoBRET signal with Cyclin E1 (**Table S13**)^58^.

## Discussion

In the field of drug discovery and development of chemical biology tools the initial hit-finding phase can often be long, tedious, and particularly prone to experimental artefacts. Although all assays suffer to some extent from these issues, a robust and generalizable hit-finding strategy that can be executed at scale will be highly enabling for many applications in medicine and chemical biology. In testing the broad applicability of DEL-ML, we applied this approach to 18 WDRs with a range of druggability^34^ scores (**Figs. 2b,c**), and two non-WDRs with ligandable PPI domains as reference points. It is worth noting, that assessing assay interference, for example, solubility and aggregation of compounds is a key part of the workflow.

To our knowledge this is the first time that a large cohort of proteins were subjected to DEL selection followed by ML prediction of ligands in a systematic and coordinated manner. This allowed us not only to gain insight as to the general applicability of the DEL-ML method, but also to make conclusions about the ligandability of the WDR family in particular. Our discovery of ligands for almost half of the tested WDR domains clearly demonstrates their ligandability, and of the promise of the DEL-ML approach in general. Moreover, computational druggability analysis using AlphaFold models of over 300 human WDR proteins suggested that ∼ 50% of WDRs are ligandable. This is a conservative estimate considering that cryptic pockets are often hidden in apo WDR structures (**Fig. 1, Tables S5, S6**). Indeed, DEL-ML selection yielded novel, selective hits for previously unliganded WDRs, of which two are predicted to be below the ‘ligandable threshold’ (**Fig. 2c, Table S7**).

With the exception of WDR5 we did not address whether our new ligands could modulate protein function, such as disruption of PPIs. However, it is quite likely that potent ligands that bind in the central pocket of the WDR donut would modulate key PPI functions. Moreover, ligands that are not able to modulate a PPI are highly valuable starting points for development of proximity-based pharmacological agents such as PROTACs. In this regard it is notable that several WDRs act as substrate binding modules of E3 ligases to recruit substrates for polyubiquitylation and degradation by the proteasome^3^. These include most of the substrate binding domains of the Cullin 4 E3 ligase family^20,59^, the FBXW class of Cullin 1 and 7 E3 ligase families such as FBXW5, FBXW7, and some non-Cullin E3 ligases such as RFWD2 and RFWD3. In addition, several other WDR proteins such as WDR5, EED and WDR12^21^ interact with DDB1, which is itself a WDR domain-containing substrate adaptor of the CUL4 E3 ligases. Therefore, new small molecule ligands that bind to WDR domain-containing E3 ligases may expand the repertoire of E3 ligases available for PROTAC development^60^.

## Outlook

Our demonstration of the broad applicability of DEL-ML as a practical strategy for ligand discovery at scale has important implications for tackling the wider human proteome to identify starting points for pharmacological modulators such as chemical probe and drugs^22^. Our systematic establishment of protein-production and crystallization protocols and our discovery of small molecule ligands for WDR containing proteins should serve as a highly enabling resource to investigate the function of WDR proteins and their potential for therapeutic targets. Indeed, the ensemble of resources presented here should be seen as an invitation to explore the unknown biology of the dark WDR-ome. We hope that this work will inspire similar efforts focused on other protein families, a vision originally formulated by the Target 2035 initiative^22-24^.

## Methods

### Informatics analysis

Using InterPro IDs: IPR001680 IPR036322 IPR015943 as the search query, a list of 349 WDR domain proteins and the number of publications and the number of articles associating each protein with specific disease areas were sourced from chembioport.thesgc.org, which extracts data from the NCBI’s gene2pubmed database (**Table S1**)^61^. The amino acid sequence of each WDR domain protein was sourced from UniProt and a multiple sequence alignment (MSA) was generated using ICM-Pro 3.9-3a (Molsoft, San Diego). The data collected and MSA was then used to generate annotated phylogenetic trees using iTOL v6.

### Druggability analysis

AlphaFold V2.0 models for each of the 349 WDR proteins (**Table S5**) were downloaded from https://alphafold.ebi.ac.uk/ ^32,62^. Structures were unavailable for eight proteins (UniProt IDs: P50851, Q5T4S7, Q6ZNJ1, Q6ZS81, Q8IZQ1, Q8NFP9, Q9H799, Q99698) and thus were excluded from the analysis. After visual inspection, six proteins were excluded from further analysis due to the lack of a complete ‘donut’-shaped domain (UniProt IDs: P18583, P55735, O75460, Q76MJ5, Q9NZJ5, Q96EE3). When multiple WDR domains were found in a protein, they were all analyzed. Each WDR domain was subjected to the ICMPocketFinder algorithm using ICM-Pro 3.9-3a (Molsoft, San Diego) to predict to predict the location of potential pockets and to obtain the drug-like density (DLID) score^34^. The domain was then superimposed with a canonical WDR structure (WDR5, PDB ID: 4QL1) with 4 dummy atoms placed down the middle of the central pocket. The DLID value of the pocket with the smallest distance to any of these 4 atoms was reported as the druggability index for the central pocket. For proteins with multiple WDR domains, the highest DLID score was reported. Proper superimposition was confirmed through visual inspection. To assess the predictive confidence of the pocket, the number of residues within 5Å from the pocket with an AlphaFold confidence score (pLDDT) < 70 were reported (**Table S5**). DLID scores for pockets where no such residues are found are expected to be more reliable.

### Biotechnology

Individual expression clones are detailed in **Table S2** and are available from Addgene. Protein expression and purification summaries are detailed in **Table S3** and deposited in Zenodo^37^.

### Structural biology

Twenty-one, high resolution structures have been deposited in the PDB (**Table S4**). The data collection and refinement statistics are in **Table S10**.

### DNA Encoded Library Selection

Affinity mediated selections included a pool of more than sixty DNA-encoded libraries**^63^**. For each His6-tagged target the DEL deck was combined with the protein and incubation occurred in solution prior to capture on the IMAC matrix, for biotinylated targets the target was captured on the streptavidin matrix prior to blocking with biotin and the subsequent introduction of the DEL deck (**Table S6**). For each target an additional selection condition containing no target was performed in parallel to identify matrix binders. For some targets an additional selection condition containing target protein with additive, e.g. a competitor, was performed in parallel (**Table S6).**

For each IMAC selection (no target, target, or target with competitor), purified protein was pre-incubated for 30 minutes with or without additive with the DEL deck (40 μM) for 1 h in a volume of 60 μL in 1× selection buffer then a separate ME200 tip (Phynexus) containing appropriate affinity matrix was prewashed three times in 200 μL of fresh 1× selection buffer. Each selection was separately captured with 20 passages over the ME200 tip over 0.5 h. The bound protein and associated library members were captured on the ME200 tip and washed eight times with 200 μL of, fresh 1× selection buffer. Bound library members were eluted by incubating the ME200 tip with 60 μL of 1× fresh selection buffer at 85 °C for 5 min. The elution was incubated with 20 passages over a fresh, prewashed ME200 tip containing appropriate affinity matrix to remove any eluted protein. The selection process was run for a second time using the eluate of the first selection in place of the input DNA-encoded library and using appropriate fresh target or no target.

For streptavidin matrix selections (no target, target, or target with competitor), purified protein was pre-incubated for 30 min with or without additive in a volume of 60 μL in 1× selection buffer. A separate ME200 tip (Phynexus) containing the affinity matrix was prewashed three times in 200 μL of fresh 1× selection buffer. Proteins were captured with 20 passages over the ME200 tip for a total of 0.5 h. The bound protein captured on the ME200 tip was washed two times with 200 μL of, fresh 1× selection buffer (with 50 μM biotin) then the DEL deck (40 μM) was introduced in a volume of 60 μL in 1× selection buffer with or without additive was passaged over the ME200tip 40 times over the period of an hour. ME200 tips were then washed eight times with 200 μL of, fresh 1× selection buffer. Bound library members were then eluted by incubating with 60 μL of 1× fresh selection buffer at 85 °C for 5 min. The elution was then subjected to 20 passages over a fresh, prewashed ME200 tip containing to remove any eluted protein. The selection process was run for a second time using the eluate of the first selection round in place of the input DNA-encoded library and using appropriate fresh target or no target.

For both IMAC and Streptavidin matrix selections, the eluate of the second round of selection was PCR amplified and sequenced^63^. Sequence data were parsed, error-containing sequences were disregarded, amplification duplicates were removed, and building block and chemical scheme encodings were decoded and reported along with associated calculated statistical parameters.

The immobilization efficiency of proteins during selection experiments was determined by SDS-PAGE. Input sample is identical to the initial solution containing protein in 60 μL 1x selection buffer prior to capture with 20 passages over the ME200 tip. Flow-through sample is the solution containing protein in 60 μL 1x selection buffer after capture with 20 passages over the ME200 tip from selection. Resin sample is the resin removed from the ME200 tip and mixed to resuspension in 60 μL 1x selection buffer. All proteins demonstrated ≥ 80% purity and ≥ 50% fraction of input protein immobilized (**Fig S5**).

#### ML Models Generated using Random Forest

For the random forest models (**Table S7**), disynthons were classified into two groups: “non-hits” (label = 0) and either “hits” or “competitive hits” (label = 1), depending on whether the selection experiment included a condition with a competitive binder. Thirty models were trained, with one million negative examples randomly sampled from the “non-hits” and 90% randomly sampled from the “hits” or “competitive hits” as positive examples to make up the training set for each model. 10% of this total set of examples was held out as a test set and was not seen during training. The positive (label = 1) examples were oversampled by including them twice during the training for each model. Each molecule in the training and test sets was featurized using the Morgan fingerprint algorithm as implemented in the RDKit cheminformatics package with radius=2 and bit count=1024^64^. The models were trained using the RandomForestClassifier class in the scikit-learn Python package^65^. Non-default hyperparameters for the models included n_estimators=2000 and min_samples_split=5. Of the thirty trained models, the model with the highest accuracy on the test set was chosen for inference on commercial catalogs.

#### ML Models Generated using GCNN

The GCNN model in this study (**Table S7**) is based on that in McCloskey et al.^25^, with several improvements referred to as GCNN V1, V2 and V2.1. We elaborate on all 3 versions below.

#### Label assignment for ML

DNA Encoded Libraries molecules were aggregated into disynthons and enrichment values were calculated following the approach detailed in McCloskey et al.^20^. Subsequently, each disynthon was assigned to one of five distinct classes: competitive hit, non-competitive hit, promiscuous binder, matrix binder, or non-hit^20^. For targets where a competitor molecule was incubated during a DEL affinity screen, the positive training examples for machine learning were designated as the "competitive hit" class. For the targets without competitor molecules, a target hit class replaced both competitive hit and non-competitive hit classes, resulting in four instead of five classes.

**GCNN V1** adopted the model from McCloskey et al.^25^ with the following updates: 1) The model training was migrated from CPU to tensor processing units (TPU) to take advantage of TPU’s strength in synchronous training. 2) In addition to ensembling the cross-validation fold models as described in McCloskey et al.^25^, we implemented two extra levels of model ensembling to improve overall reliability: a) We used TPU-supported graph partitioning for manual model parallelism, training 8 models with independently randomized weight initialization on individual TPU cores. The ensembled prediction is the median of prediction scores from these 8 models. b) We further replicated the full training process 3 times, and the final prediction score is the median of predictions from the 3 runs, each with 8 TPU model replicas, and the cross-validation fold models.

**GCNN V2** built upon GCNN V1 with the following key improvements: 1) Instead of using the maximum ROC-AUC we utilized the top_100_actives metric. This metric counts the number of active examples within the top 100 ranked predictions for the hit class. After training models for each cross-validation fold, we selected the checkpoint weights with the highest top_100_actives score on the tuning set. 2) Along with changing the model selection metric, we also updated training hyperparameters: a) Learning rate was increased from 0.0006 to 0.1. b) Training steps were increased from 30,000 to 1,000,000, with checkpoints saved every 100,000 steps instead of every 3,000 steps.

**GCNN V2.1** is an ensemble model combining the predictions from GCNN V2 and a molecular fingerprint-based deep neural network (DNN). The fingerprint DNN architecture consists of 2 fully connected layers of sizes (2000, 100) with rectified linear activation function (ReLU) and the input features are 2048-bit Morgan Fingerprint with radius of 3 (ECFP6_2048). The ensemble procedure involves first retrieving the top 100,000 compounds using the GCNN V2 model predictions, and then filtering out 50% of these compounds based on the lower DNN prediction scores, and from the remaining 50,000 compounds, following the same protocol of property filtering and diversity selection, ranked by GCNN V2 predictions, to select the compounds for purchase and testing.

#### ML Models Generated using ArtemisAI

Where the model type is specified as “ArtemisIAI” (**Table S7**), the DEL selection data were featurized in a range of different formats and each format was then used to build a collection of models. These models were evaluated and, based on performance, included in a model ensemble used to score commercial catalogs compounds. A diverse set of the highest scoring compounds were ordered and assayed.

#### DEL-ML-driven hit expansion

For promising hit compounds like those found for WDR12, DCAF1, and WDR91, we performed automated hit expansion using the GCNN model to search for and prioritize analogs of the initial hit. The search was performed in the large Enamine REAL library of 1.9 billion commercially available compounds, avoiding expensive bespoke synthesis. First, we identified analogs from the Enamine REAL library with ECFP6 Tanimoto similarity above a threshold to the original hit. We then used the GCNN prediction score to rank these compounds. To balance exploiting structural similarity and exploring diversity, we also applied DISE selection with a 0.2 radius cut-off to increase analog diversity.

### Computationally based hit expansion

The EnamineReal commercially available structural space most similar to the 4 initial DCAF1 hits (Tanimoto Score >0.35 using ECFP6, ∼87 thousand compounds) was docked (ICM and OpenEye ORION platform) in the X-Ray structure and compounds with high docking scores were ordered and assayed. A similarly set was also selected from the highest docking scores (ICM and OpenEye ORION platform) from a 10M random sampling of EnamineReal. Lastly, a 3D shape-based ROCS hit expansion search (Openeye ORION platform) of the 4 original hits was conducted and a set of compounds with good shape complementarity (Tanimoto combo score > 1.4) to the original hits were ordering and assay.

### Surface plasmon resonance (SPR)

The binding affinity of compounds was assessed by SPR (Biacore™ T200 or Biacore™ 8K, Cytiva Inc.) as previously described^66^. SPR running conditions (**Table S9**) vary slightly depending on the specific compounds and protein targets.

### Differential Scanning Fluorimetry (DSF)

WDR5 was diluted to 0.1 mg/mL in buffer (100 mM HEPES, 100 mM NaCl, pH 7.5) in the presence of 5 x SYPRO Orange dye (Life Technologies, S-6650) and serially titrated MR44397 (up to 200 µM) in a total volume of 20 µl in a white polypropylene 384-well plate (Axygen, #UC500). DSF was performed in a Light Cycler 480 II (Roche Applied Science, Penzberg, Germany) using a 4 °C/min temperature gradient from 20 °C to 95 °C. Data points were collected at 1 °C intervals. DSF data was fitted to a Boltzmann sigmoid function and T_m_ values were determined as previously described^6^.

### Fluorescence Polarization (FP) Assay

WDR5 (H3 1-15 FP: 300 nM, WIN and RBBP5 FP: 5 µM) was incubated with 40 nM of fluorescent FITC labelled peptide- H3 1-15, -WIN (GSARAEVHLRKS) or - RBBP5 (371-380aa, EDEEVDVTSV) in the presence or absence of serial titration of MR44397 (up to 100 µM) for 30 min at ambient temperature, and fluorescence polarization was measured as described before^67^.

SETDB1 TTD (197-403:) 5 µM was incubated with 40 nM of 5’ FITC-H3K9me2K14ac (1-25) with serially diluted MR46747 compound at 200 μM top concentration (12 points 3-fold dilution). Final DMSO in the reaction mixture was maintained at 2%. Following a brief spin down and incubation for 30 min at room temperature fluorescence polarization ratio measurements were obtained as reported previously^67^.

### Bio-layer Interferometry (BLI) assay

The BLI assay was used to evaluate the binding kinetics of MR44397 against WDR5. All assays were run on the same Octet Red96 instrument at 25 °C with shaking at 1000 rpm. BLI assay buffer consists of 0.1% BSA in (10 mM HEPES, pH 7.4, 150 mM NaCl, 0.05% Tween 20, 3 mM EDTA). Anti-biotin super streptavidin (SSA) biosensors (part no: 18-5057, FORTÉBIO, Fremont, California, United States) were loaded into the columns of a biosensor holding plate and pre-hydrated in BLI assay buffer for 10 min. Black 96-well microplates were loaded with 200 μL per well. Following a 60-s equilibration step with anti-biotin super streptavidin (SSA) biosensors in buffer, biotinylated WDR5 was loaded for 300 s. The SSA were then moved into wells in the following sequence: (i) a baseline step with fresh assay buffer for 60 s, (ii) MR44397 for a 60 s association step, and (iii) assay buffer for a 120 s dissociation step. Binding curves from the association and dissociation steps were fitted using a 1:1 binding model to determine K_D_ values using the Octet Data Acquisition software.

### Dynamic Light Scattering (DLS)

The solubility of compounds was estimated by DLS that directly measures compound aggregates and laser power in solution. Compounds were serially diluted directly from DMSO stocks, then diluted 50x into filtered HBS-E (20 mM HEPES pH 7.4, 150 mM NaCl, 3 mM EDTA) for analysis by DLS (DynaPro DLS Plate Reader III, Wyatt Technology, USA) as previously described^68^. The solubility is tabulated alongside binding data in Table S8.

### ^19^F-NMR spectroscopy

The binding of fluorinated compounds was assayed by looking for the broadening and/or perturbation of ^19^F resonances upon addition of LRKK2 (at protein to compound ratios of 0.5:1 to 4:1) in PBS buffer (pH7.4, 137 mM NaCl, 2.7 mM KCl, 10 mM Na_2_HPO_4_, 1.8 mM KH_2_PO_4_, and with 5% D_2_O). 1D-^19^F spectra were collected at 298K on a Bruker AvanceIII spectrometer, operating at 600 MHz, and equipped with a QCI probe. Two to four thousand transients were collected with an acquisition period of 0.2 s, over a sweep width of 150 ppm, a relaxation delay of 1.5 s, and using 90° pulses centered at -120 ppm. The concentration of the compounds in both reference and protein-compound mixtures was 5-10 µM. TFA (20 µM) was added as an internal standard for referencing. Prior to Fourier transformation, an exponential window function was applied (lb = 1 to 3) to the FID. All processing was performed at the workstation using the software Topspin3.5.

### Hydrogen-Deuterium Exchange Mass Spectrometry

HDX-MS was performed as previously described using the PAL3 Autosampler (CTC Analytics AG) coupled to an M-Class UPLC (Waters Corp., U.K.) and SELECT SERIES Cyclic IMS Mass Spectrometer (Waters Corp., U.K.) in HDMSe mode^69^. Briefly, samples (5 μM WDR12 or 5 μM WDR12 + 100 μM MR44915) were labeled (10 mM Phosphate Buffer pD 7.4, 150 mM NaCl) at 20 °C for 0, 1, 10, 30, and 60 mins. The HDX reaction was then quenched (100 mM Phosphate Buffer, pH 2.5) at 0 °C for 1 min prior to proteolysis (Enzymate BEH Pepsin column, Waters Corp., U.K.), desalting (ACQUITY UPLC BEH C18 VanGuard Pre-column, Waters Corp., U.K.), and reverse-phase separation (ACQUITY UPLC BEH C18 Column, Waters Corp., U.K.). Data processing was conducted using ProteinLynx Global Server 3.0.3. and DynamX 3.0 (Waters Corp., U.K.), and visualization was done using PyMol 2.5.0 (Schrodinger, LLC). For a cumulative ΔHDX-MS peptide signal to be considered statistically significant, it must have exceeded the cumulative propagated error (2 σ, n ≥ 2).

### NanoBRET-based Protein-Protein Interaction (PPI)

A detailed protocol was recently published^29^. Briefly, HEK293T cells were seeded in 96-well white plates (4 x 10^4^ cells/well, Corning) and co-transfected with C-terminally NL-tagged WDR5 (0.001 µg/well) and C-terminally HT-tagged histone H3 (0.1 µg/well) or N-terminally NL-tagged WDR5 (0.001 µg/well), C-terminally HT-tagged RBBP5 (0.03 µg/well) and empty vector (0.07 µg/well) using XtremeGene HP transfection reagent, following manufacturers’ instructions. The next day the media was replaced with DMEM (no phenol red) supplemented with 4% FBS, 100 U/mL penicillin, and 100 μg/mL streptomycin, +/- 1000-fold diluted HaloTag^®^ NanoBRET™ 618 Ligand (Promega) and incubated with indicated compounds concentrations or DMSO control for 4 h. Next, 100-fold diluted NanoBRET™ Nano-Glo^®^ Substrate (Promega) was added and donor emission at 450 nm and acceptor emission at 618 nm were read within 10 min of substrate addition using a ClarioStar plate reader (Mandel Scientific). The NanoBRET ratio was determined by subtracting 618/460 (acceptor/donor) signal from cells without NanoBRET™ 618 HaloTag Ligand x 1000 from 618/460 signal from cells with NanoBRET™ 618 Ligand x 1000.

### Cellular interaction between small molecules and proteins

#### NanoLuciferase-based Cellular Thermal Shift Assay

A protocol was recently published^28^. Briefly, HEK293T cells were seeded in 6-well white plates (8 x 10^5^ cells/well, Corning) and transfected with N-terminally HiBiT-tagged WDR5 (0.2 µg/well) and empty vector (1.8 µg/well) using XtremeGene HP transfection reagent, following manufacturers’ instructions. The next day cells were trypsinized and resuspended in optiMEM (no phenol red, Gibco) at the density of 2 x 10^5^ cells/mL. After compound addition (50 µM of MR44397 and 20 µM of OICR-9429) or DMSO, the cells were aliquoted in a 96-well PCR plate (one well per intended temperature, 50 µl), covered with breathable film and incubated for 1 h at 37 °C 5% CO_2_. Next, cells were heated in the VerityPro thermocycler (Thermo Fisher Scientific) as follows: 1 min at (22 °C) and 3 min at the indicated temperature gradient, followed by a cooling step to chill samples to RT (22°C). Next, 50 µl of lytic buffer was added (2% NP-40, protease inhibitors (Roche), 0.2 µM LargeBiT (produced in house) in optiMEM (no phenol red)). After 10 min incubation at RT, 25 µl/well of 100-fold diluted NanoBRET™ Nano-Glo^®^ Substrate (Promega) was added and transferred to 384-well white plates in quadruplicates (20 µl/well). The luminescence signal was read in the ClarioStar plate reader (Mandel Scientific). Data were fitted to obtain apparent *T*_agg_ values using the nonlinear curve fit (e.g., Boltzmann Sigmoid equation) using GraphPad Prism software.

#### Tracer-based NanoBRET assay

HEK293T cells were plated in a 6-well plate (8 x 10^5^ cells/well) and transfected with 0.2 µg of N-terminally NL-tagged WDR5 vector and 1.8 µg of empty vector using XtremeGene HP transfection reagent, following manufacturer’s instructions. The next day cells were trypsinized and resuspended in optiMEM (no phenol red) at 2 x 10^5^ cells/mL density with 1 µM of WDR5 tracer^54^. Compound serial dilutions were prepared in DMSO and added to cells. Cells were transferred to 384-well low-binding white plates (20 µl/well, Corning) in quadruplicates. After 2 h, 5 µl/well NanoBRET™ Nano-Glo® Substrate (Promega) and Extracellular NanoLuc^®^ Inhibitor (Promega) diluted in optiMEM (no phenol red) 100-fold and 300-fold, respectively, was added. The donor emission at 450 nm and acceptor emission at 618 nm were read immediately after substrate addition and shaking plate for 20 s using a ClarioStar plate reader (Mandel Scientific). NanoBRET ratios were calculated by subtracting the mean of 610 nm/460 nm signal from cells without tracer x 1000 from the 610 nm/460 nm signal from cells with tracer x 1000. Data were fitted to obtain IC_50_ values using the nonlinear curve fit using GraphPad Prism software.

### Experimental Procedure of 1-acetyl-N-((2-(2-chlorophenoxy)pyridin-4-yl)methyl)indoline-5-sulfonamide MR44915

#### Reaction Scheme

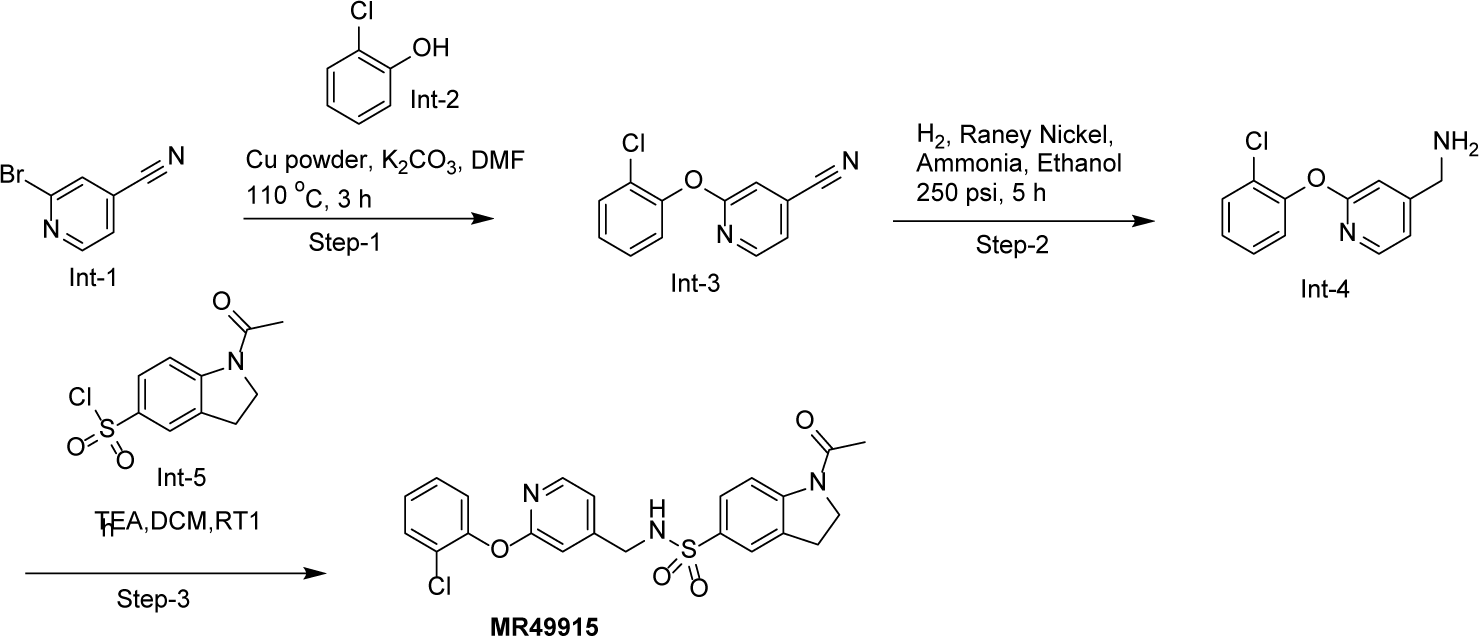

#### Step-1: Synthesis of 2-(2-chlorophenoxy) isonicotinonitrile (Int-3)

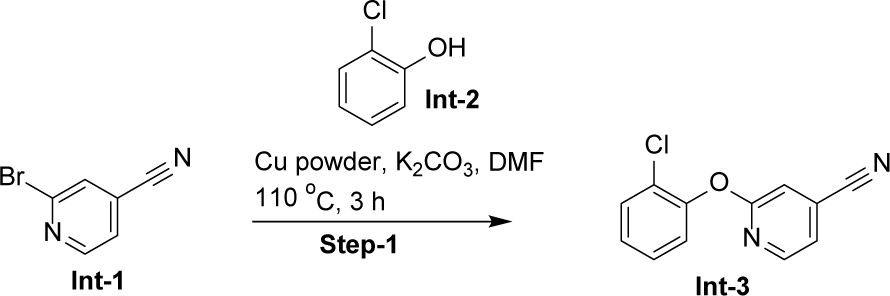

To a mixture of 2-bromoisonicotinonitrile **(Int-1)** (0.2 g, 1.09 mmol, 1.0 eq.) and 2-chlorophenol **(Int-2)** (0.2 g, 1.63 mmol, 1.5 eq.) in *N, N*-dimethylformamide (2 mL), were added potassium carbonate (0.23 g, 1.63 mmol, 1.5 eq.) and copper powder (0.04 g, 0.655 mmol, 0.2 eq.), and the mixture was stirred for 3 h at 110 °C. The reaction mixture was poured into water and extracted with ethyl acetate dried over sodium sulphate and concentrated under reduced pressure to give the crude compound, which was purified by column chromatography (SiO_2_, ethyl acetate: hexanes = 1:10 to 2:10) to give 2-(2-chlorophenoxy) isonicotinonitrile **(Int-3)** (0.17 g, yield: 67.63 %) as a brown solid.

**LC-MS**: m/z=232.8 [M+2H]^2+^.

#### Step-2: Synthesis of (2-(2-chlorophenoxy) pyridin-4-yl) methanamine (Int-4)

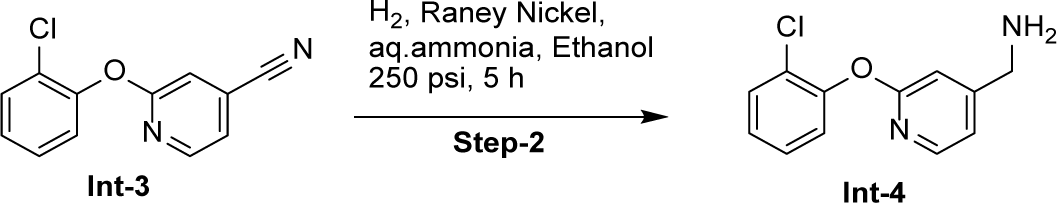

To a solution of 2-(2-chlorophenoxy) isonicotinonitrile (**Int-3**) (0.17 g, 0.737 mmol, 1.0 eq.) in ethanol (11.2 mL) in autoclave under nitrogen atmosphere, aqueous ammonia (1.1 mL) and Raney nickel (0.2 g) were added and stirred under 250 psi pressure of hydrogen for 5 h. The reaction mixture was filtered through a celite pad and the filtrate was concentrated under reduce pressure to give crude product (2-(2-chlorophenoxy) pyridin-4-yl) methanamine **(Int-4)** (0.16 g, yield: 92.50 %) as a brown solid, which was used in the next step without further purification.

**LC-MS**: m/z=235.0 [M+H]^+^.

#### Step-3: Synthesis of 1-acetyl-N-((2-(2-chlorophenoxy)pyridin-4-yl)methyl)indoline-5-sulfonamide MR44915

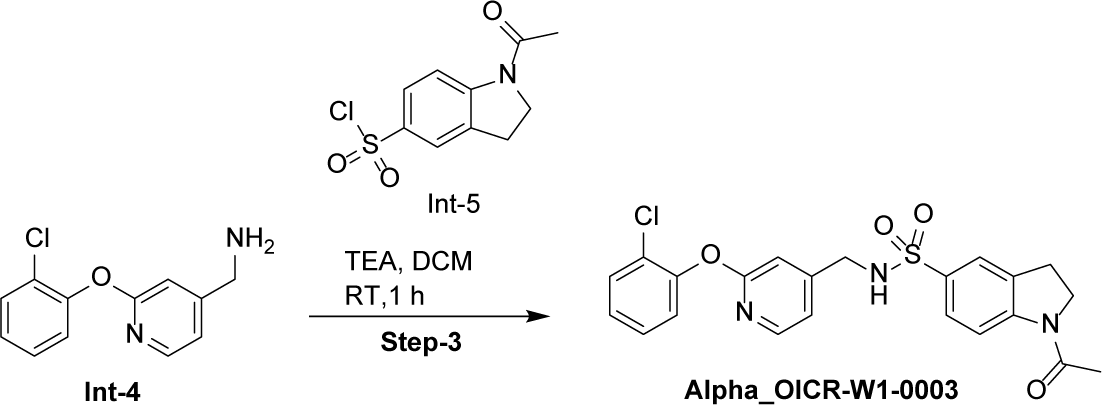

To a cooled to solution (0 °C) of (2-(2-chlorophenoxy) pyridin-4-yl) methanamine **(Int-4)** (0.16 g, 0.68 mmol, 1.0 eq.) in dichloromethane (3.2 mL), triethylamine (0.2 g, 2.04 mmol, 3.0 eq.) and 1-acetylindoline-5-sulfonyl chloride **(Int-5)** (0.23 g, 0.88 mmol, 1.3 eq.) were added. The mixture was then stirred for 1 h at room temperature. The reaction mixture was poured into water and extracted with dichloromethane, dried over sodium sulphate and concentrated under reduced pressure to give the crude compound which was purified by column chromatography using (SiO_2_, ethyl acetate: hexanes = 2:10 to 4:10) to give 1-acetyl-N-((2-(2-chlorophenoxy) pyridin-4-yl) methyl) indoline-5-sulfonamide **(MR44915)** (0.07 g, yield: 22.40 %) as a white solid.

**LC-MS:** m/z = 458.0[M+H]^+^.

**^1^H NMR (400 MHz, DMSO d_6_):** δ 8.23 (s, 1H), 8.16 (d, J = 8.8 Hz, 1H), 7.99 (d, J = 5.2 Hz, 1H), 7.62 – 7.39 (m, 3H), 7.41 (t, J = 7.8 Hz, 1H), 7.31– 7.23 (m, 2H), 7.04 (d, J = 5.3 Hz, 1H), 6.97 (s, 1H), 4.17 (t, J = 8.7 Hz, 2H), 4.09 (d, J = 6.3 Hz, 2H), 3.19 (t, J = 6.5 Hz, 2H), 2.21 (s, 3H).

**^13^C NMR (101 MHz, DMSO d_6_):** δ 170.22, 163.04, 152.23, 149.77, 146.71, 134.91, 133.58, 130.81, 128.97, 127.03, 124.74, 123.77, 118.49, 115.75, 109.41, 49.16, 45.25, 27.52, 24.60

## Supporting information

Supplementary Figures and short tables

## Data and Reagent Availability

Experimental 3D protein structure coordinates are deposited in the Protein Data Bank under accession codes listed in **Table 10**. Protein expression clones have been deposited to Addgene (and are listed in **Table S2** as reference). Purification protocols and associated data for all WDR proteins manufactured within this study are available at DOI: 10.5281/zenodo.10655440.

## Abbreviations

BLI: Biolayer Interferometry
DEL: DNA Encoded Library
DLID: drug-like density
DSF: Differential Scanning Fluorimetry
FP: Fluorescence Polarization
GCNN: Graph Convolutional Neural Network
HDX: Hydrogen-Deuterium eXchange
HTS: High-Throughput Screening
ML: Machine Learning
NanoBRET: NanoLuciferase Bioluminescence Resonance Energy Transfer
PPI: Protein-Protein Interaction
PROTAC: Proteolysis Targeting Chimera
SPR: Surface Plasmon Resonance
T_m_: Protein melting temperature
WDR: Tryptophan-Aspartate Repeat

## Acknowledgements

The authors acknowledge Dr Brian Wilson for determining the purity of the compounds presented in the main body of the manuscript, and Dr Drashti Thumar for synthesizing the chlorine analog MR44915. The Structural Genomics Consortium is a registered charity (no: 1097737) that receives funds from Bayer AG, Boehringer Ingelheim, Bristol Myers Squibb, Genentech, Genome Canada through Ontario Genomics Institute [OGI-196], EU/EFPIA/OICR/McGill/KTH/Diamond Innovative Medicines Initiative 2 Joint Undertaking [EUbOPEN grant 875510], Janssen, Merck KGaA (aka EMD in Canada and US), Pfizer, and Takeda.

We thank the staff at the Northeastern Collaborative Access Team, which is funded by the National Institute of General Medical Sciences from the National Institutes of Health (P30 GM124165). The Eiger 16M detector on the 24-ID-E beamline is funded by an NIH-ORIP HEI grant (S10OD021527). This research used resources of the Advanced Photon Source, a U.S. Department of Energy (DOE) Office of Science user facility operated for the DOE Office of Science by Argonne National Laboratory under contract no. DE-AC02-06CH11357.

## Author Contributions

AME, CHA: Concept, Leadership, and Funding acquisition; AS, AH, EG, HZ, MK, MSilva, MYPG, PL, YL: Protein manufacturing; ADong, GLR, LH, SAhmad, SB, SK, SS: Structural biology; MvR, JPG, JWC, RDT: Selection Design and Analysis; ADK, JPG, MAC, PAC, RDT, YZ: DNA-Encoded Chemical Library Synthesis and Selection; ADumas, AJB, BG, BLS, CJM, CM, CU, ES, IW, JAF, JD, JG, JL, JT, JX led GCNN model application & development, JZ, MAG, NH, PR, SK, TK, WT, YA, YZ: Machine-Learning and Compound Design; AB, AKY, AL, CL, DJW, EW, FL, GLR, HAK, IC, JL, MDS, MSilva, OH, PJB, SAckloo, SAhmad, SB, SDT, SG, SH, SP, VS: Biophysical assays; MSzewczyk, RACM: Cell Assays; PJB, SAckloo: Project Management; Supervision: ADK, AME, CHA, DB-L, JAF, JWC, LH, MAC, MAG, MSchapira, PAC, YZ; CHA, and All: Writing, Reviewing, Editing.

## References

1. Song, R., Wang, Z.D. & Schapira, M. Disease Association and Druggability of WD40 Repeat Proteins. Journal of proteome research 16, 3766–3773 (2017).

2. Wang, J., Yazdani, S., Han, A. & Schapira, M. Structure-based view of the druggable genome. Drug Discov Today 25, 561–567 (2020).

3. Schapira, M., Tyers, M., Torrent, M. & Arrowsmith, C.H. WD40 repeat domain proteins: a novel target class? Nature reviews.Drug discovery 16, 773–786 (2017).

4. Xu, C. & Min, J. Structure and function of WD40 domain proteins. Protein Cell 2, 202–214 (2011).

5. Yeh, J.I., et al. Damaged DNA induced UV-damaged DNA-binding protein (UV-DDB) dimerization and its roles in chromatinized DNA repair. Proc Natl Acad Sci U S A 109, E2737–2746 (2012).

6. Grebien, F., et al. Pharmacological targeting of the Wdr5-MLL interaction in C/EBPalpha N-terminal leukemia. Nature chemical biology 11, 571–578 (2015).

7. Getlik, M., et al. Structure-Based Optimization of a Small Molecule Antagonist of the Interaction Between WD Repeat-Containing Protein 5 (WDR5) and Mixed-Lineage Leukemia 1 (MLL1). J Med Chem 59, 2478–2496 (2016).

8. Teuscher, K.B., et al. Structure-based discovery of potent WD repeat domain 5 inhibitors that demonstrate efficacy and safety in preclinical animal models. Proc Natl Acad Sci U S A 120, e2211297120 (2023).

9. Senisterra, G., et al. Small-molecule inhibition of MLL activity by disruption of its interaction with WDR5. Biochem J 449, 151–159 (2013).

10. He, Y., et al. The EED protein-protein interaction inhibitor A-395 inactivates the PRC2 complex. Nat Chem Biol 13, 389–395 (2017).

11. Qi, W., et al. An allosteric PRC2 inhibitor targeting the H3K27me3 binding pocket of EED. Nat Chem Biol 13, 381–388 (2017).

12. Simonetta, K.R., et al. Prospective discovery of small molecule enhancers of an E3 ligase-substrate interaction. Nat Commun 10, 1402 (2019).

13. Sackton, K.L., et al. Synergistic blockade of mitotic exit by two chemical inhibitors of the APC/C. Nature 514, 646–649 (2014).

14. Li, A.S.M., et al. Discovery of Nanomolar DCAF1 Small Molecule Ligands. J Med Chem (2023).

15. Tao, Y., et al. Targeted Protein Degradation by Electrophilic PROTACs that Stereoselectively and Site-Specifically Engage DCAF1. J Am Chem Soc 144, 18688–18699 (2022).

16. Vulpetti, A., et al. Discovery of New Binders for DCAF1, an Emerging Ligase Target in the Targeted Protein Degradation Field. ACS Med Chem Lett 14, 949–954 (2023).

17. Schroder, M., et al. DCAF1-based PROTACs with activity against clinically validated targets overcoming intrinsic- and acquired-degrader resistance. Nat Commun 15, 275 (2024).

18. Zhang, X., et al. DCAF11 Supports Targeted Protein Degradation by Electrophilic Proteolysis-Targeting Chimeras. J Am Chem Soc 143, 5141–5149 (2021).

19. Li, Y.D., et al. Template-assisted covalent modification of DCAF16 underlies activity of BRD4 molecular glue degraders. bioRxiv (2023).

20. Higa, L.A., et al. CUL4-DDB1 ubiquitin ligase interacts with multiple WD40-repeat proteins and regulates histone methylation. Nat Cell Biol 8, 1277–1283 (2006).

21. Lee, J. & Zhou, P. DCAFs, the missing link of the CUL4-DDB1 ubiquitin ligase. Mol Cell 26, 775–780 (2007).

22. Carter, A.J., et al. Target 2035: probing the human proteome. Drug discovery today 24, 2111–2115 (2019).

23. Müller, S., et al. Target 2035 - update on the quest for a probe for every protein. RSC Med Chem 13, 13–21 (2022).

24. Ackloo, S., et al. Target 2035 – an update on private sector contributions. RSC Med Chem (2023).

25. McCloskey, K., et al. Machine Learning on DNA-Encoded Libraries: A New Paradigm for Hit Finding. J Med Chem 63, 8857–8866 (2020).

26. Torng, W., et al. Deep Learning Approach for the Discovery of Tumor-Targeting Small Organic Ligands from DNA-Encoded Chemical Libraries. ACS Omega 8, 25090–25100 (2023).

27. Ahmad, S., et al. Discovery of a First-in-Class Small-Molecule Ligand for WDR91 Using DNA-Encoded Chemical Library Selection Followed by Machine Learning. J Med Chem 66, 16051–16061 (2023).

28. Ramachandran, S., et al. HiBiT Cellular Thermal Shift Assay (HiBiT CETSA). Methods Mol Biol 2706, 149–165 (2023).

29. Szewczyk, M.M., Owens, D.D.G. & Barsyte-Lovejoy, D. Measuring Protein-Protein Interactions in Cells using Nanoluciferase Bioluminescence Resonance Energy Transfer (NanoBRET) Assay. Methods Mol Biol 2706, 137–148 (2023).

30. Hui, R. & Edwards, A. High-throughput protein crystallization. J Struct Biol 142, 154–161 (2003).

31. Tunyasuvunakool, K., et al. Highly accurate protein structure prediction for the human proteome. Nature 596, 590–596 (2021).

32. Jumper, J., et al. Highly accurate protein structure prediction with AlphaFold. Nature 596, 583–589 (2021).

33. Senior, A.W., et al. Improved protein structure prediction using potentials from deep learning. Nature 577, 706–710 (2020).

34. Sheridan, R.P., Maiorov, V.N., Holloway, M.K., Cornell, W.D. & Gao, Y.D. Drug-like density: a method of quantifying the "bindability" of a protein target based on a very large set of pockets and drug-like ligands from the Protein Data Bank. J Chem Inf Model 50, 2029–2040 (2010).

35. Gurung, R., Om, D., Pun, R., Hyun, S. & Shin, D. Recent Progress in Modulation of WD40-Repeat Domain 5 Protein (WDR5): Inhibitors and Degraders. Cancers (Basel) 15(2023).

36. Wang, M., et al. Common genetic variation in ETV6 is associated with colorectal cancer susceptibility. Nat Commun 7, 11478 (2016).

37. Halabelian, L., Seitova, A., Loppnau, P., Zeng, H., Kwak, H., Li, F., Ahmad, S., Beldar, S., Bolotokova, A., Chau, I., Dehghani-Tafti, S., Dong, A., Ghiabi, P., Ackloo, S., Green, S., Herasymenko, O., Houliston, S., Hutchinson, A., Kimani, S., … Arrowsmith, C. WDR protein purification and druggability assessment. (2024).

38. Ortiz, G., Kutateladze, T.G. & Fujimori, D.G. Chemical tools targeting readers of lysine methylation. Curr Opin Chem Biol 74, 102286 (2023).

39. Guo, Y., et al. Structure-Guided Discovery of a Potent and Selective Cell-Active Inhibitor of SETDB1 Tudor Domain. Angew Chem Int Ed Engl 60, 8760–8765 (2021).

40. Mader, P., et al. Identification and characterization of the first fragment hits for SETDB1 Tudor domain. Bioorg Med Chem 27, 3866–3878 (2019).

41. Auld, D.S., Inglese, J. & Dahlin, J.L. Assay Interference by Aggregation. in Assay Guidance Manual (eds. Markossian, S., et al.) (Bethesda (MD), 2004).

42. Coussens, N.P., Auld, D.S., Thielman, J.R., Wagner, B.K. & Dahlin, J.L. Addressing Compound Reactivity and Aggregation Assay Interferences: Case Studies of Biochemical High-Throughput Screening Campaigns Benefiting from the National Institutes of Health Assay Guidance Manual Guidelines. SLAS Discov 26, 1280–1290 (2021).

43. Chan, L.L., et al. A Method for Identifying Small-Molecule Aggregators Using Photonic Crystal Biosensor Microplates. JALA Charlottesv Va 14, 348–359 (2009).

44. Schuetz, A., et al. Structural basis for molecular recognition and presentation of histone H3 by WDR5. EMBO J 25, 4245–4252 (2006).

45. Odho, Z., Southall, S.M. & Wilson, J.R. Characterization of a novel WDR5-binding site that recruits RbBP5 through a conserved motif to enhance methylation of histone H3 lysine 4 by mixed lineage leukemia protein-1. J Biol Chem 285, 32967–32976 (2010).

46. Kimani, S.W., et al. Discovery of a Novel DCAF1 Ligand Using a Drug-Target Interaction Prediction Model: Generalizing Machine Learning to New Drug Targets. J Chem Inf Model 63, 4070–4078 (2023).

47. Chow, V., Wolf, E., Lento, C. & Wilson, D.J. Developments in rapid hydrogen-deuterium exchange methods. Essays Biochem 67, 165–174 (2023).

48. Jurkowska, R.Z., et al. H3K14ac is linked to methylation of H3K9 by the triple Tudor domain of SETDB1. Nat Commun 8, 2057 (2017).

49. Vu, V., Szewczyk, M.M., Nie, D.Y., Arrowsmith, C.H. & Barsyte-Lovejoy, D. Validating Small Molecule Chemical Probes for Biological Discovery. Annu Rev Biochem 91, 61–87 (2022).

50. Martinez, N.J., et al. A widely-applicable high-throughput cellular thermal shift assay (CETSA) using split Nano Luciferase. Sci Rep 8, 9472 (2018).

51. Sanchez, T.W., et al. High-Throughput Detection of Ligand-Protein Binding Using a SplitLuc Cellular Thermal Shift Assay. Methods Mol Biol 2365, 21–41 (2021).

52. Jarzab, A., et al. Meltome atlas-thermal proteome stability across the tree of life. Nat Methods 17, 495–503 (2020).

53. Robers, M.B., et al. Target engagement and drug residence time can be observed in living cells with BRET. Nat Commun 6, 10091 (2015).

54. Dolle, A., et al. Design, Synthesis, and Evaluation of WD-Repeat-Containing Protein 5 (WDR5) Degraders. J Med Chem 64, 10682–10710 (2021).

55. Machleidt, T., et al. NanoBRET--A Novel BRET Platform for the Analysis of Protein-Protein Interactions. ACS Chem Biol 10, 1797–1804 (2015).

56. Dale, N.C., Johnstone, E.K.M., White, C.W. & Pfleger, K.D.G. NanoBRET: The Bright Future of Proximity-Based Assays. Front Bioeng Biotechnol 7, 56 (2019).

57. Castro-Castro, A., et al. Coronin 1A promotes a cytoskeletal-based feedback loop that facilitates Rac1 translocation and activation. EMBO J 30, 3913–3927 (2011).

58. Hao, B., Oehlmann, S., Sowa, M.E., Harper, J.W. & Pavletich, N.P. Structure of a Fbw7-Skp1-cyclin E complex: multisite-phosphorylated substrate recognition by SCF ubiquitin ligases. Mol Cell 26, 131–143 (2007).

59. Fouad, S., Wells, O.S., Hill, M.A. & D’Angiolella, V. Cullin Ring Ubiquitin Ligases (CRLs) in Cancer: Responses to Ionizing Radiation (IR) Treatment. Front Physiol 10, 1144 (2019).

60. Schapira, M., Calabrese, M.F., Bullock, A.N. & Crews, C.M. Targeted protein degradation: expanding the toolbox. Nature reviews.Drug discovery 18, 949–963 (2019).

61. Liu, L., Rovers, E. & Schapira, M. ChemBioPort: an online portal to navigate the structure, function and chemical inhibition of the human proteome. Database (Oxford) 2022(2022).

62. Varadi, M., et al. AlphaFold Protein Structure Database: massively expanding the structural coverage of protein-sequence space with high-accuracy models. Nucleic Acids Res 50, D439–D444 (2022).

63. Cuozzo, J.W., et al. Discovery of a Potent BTK Inhibitor with a Novel Binding Mode by Using Parallel Selections with a DNA-Encoded Chemical Library. Chembiochem 18, 864–871 (2017).

64. RDKit: Open-source cheminformatics.

65. Pedregosa, F., et al. Scikit-learn: Machine learning in Python. the Journal of machine Learning research 12, 2825–2830 (2011).

66. Dilworth, D., et al. A chemical probe targeting the PWWP domain alters NSD2 nucleolar localization. Nat Chem Biol 18, 56–63 (2022).

67. Blazer, L.L., et al. A Suite of Biochemical Assays for Screening RNA Methyltransferase BCDIN3D. SLAS Discov 22, 32–39 (2017).

68. Allen, S.J., Dower, C.M., Liu, A.X. & Lumb, K.J. Detection of Small-Molecule Aggregation with High-Throughput Microplate Biophysical Methods. Curr Protoc Chem Biol 12, e78 (2020).

69. Mann, M.K., et al. Small Molecule Screen Identifies Non-catalytic USP3 Chemical Handle. ACS Omega (2023).

