## Supplementary Figures and short tables for "A resource to enable chemical biology and drug discovery of WDR Proteins"

**Table S5. List of high-resolution 3D structures of WDR domain proteins**

| Description | PDB ID | Comment |
|---|---|---|
| WDR12 | 6N31 | Apo |
| COPA | 6PBG | Apo |
| WDR91 | 6VYC | Apo |
| WDR41 | 8W3V | Apo |
| KIF21A | 7KLJ | Apo |
| WDR55 | 7KQQ | Apo |
| CORO6 | 7KYX | Apo |
| RBBP7 | 7M3X | Apo |
| PAFAH1B1 | 7MT1 | Apo |
| UTP15 | 7RUO | Apo |
| CORO1C | 7STY | Apo |
| SEC31A | 7SUL | Apo |
| DCAF1 in complex with OICR-6766 | 7UFV | with ligand |
| COPB2 WD-domains | 8D30 | Apo |
| COPB2 WD-domain in complex with OICR-6254 | 8D41 | with ligand |
| eIF2A | 8DYS | Apo |
| DCAF1 in complex with OICR-8268 | 8F8E | with ligand |
| WDR91 in complex with MR45279 | 8SHJ | with ligand |
| WDR5 in complex with MR44397 | 8T51 | with ligand |
| WDR91 in complex with MR46654 | 8T55 | with ligand |
| SETDB1 Tandem Tudor domain in complex with MR46747 | 8UWP | with ligand |

**Table S6a. Data collection and refinement statistics**

| | CORO6 | WDR55 | WDR41 |
|---|---|---|---|
| PDB ID | 7KYX | 7KQQ | 8W3V |
| Wavelength (Å ) | 0.97918 | 0.97911 | 0.97911 |
| Resolution range (Å) | 50.00-1.63 (1.66-1.63) * | 48.75-1.80 (1.84-1.80) * | 35.0-2.20(2.24-2.20) * |
| Space group | $P2_12_12_1$ | $P2_1$ | C2 |
| Unit cell (Å) | 50.5, 80.8, 96.4 | 79.0, 58.6, 88.2 | 229.1, 86.0,52.6 |
| Total reflections | 454557 | 234127 | 219586 |
| Unique reflections | 50011(2387) | 72091(4411) | 49100(1891) |
| Multiplicity | 9.1(7.4) | 3.2(3.4) | 4.5(3.7) |
| Completeness (%) | 99.5(97.4) | 96.9 (99.5) | 95.3(74.6) |
| Mean I/sigma(I) | 24.8(1.73) | 8.2(1.4) | 22.1(1.4) |
| Wilson B-factor | 15.1 | 29.9 | 49.2 |
| R-merge | 0.092(0.942) | 0.071(0.567) | 0.066(0.953) |
| R-meas | 0.098(1.012) | 0.085(0.673) | 0.075(1.094) |
| R-pim | 0.032(0.360) | 0.045(0.365) | 0.034(0.532) |
| CC1/2 | 0.997(0.790) | 0.995(0.705) | 0.994(0.692) |
| Reflections used in refinement | 48957 | 68592 | 46626 |
| Reflections used for R-free | 989 | 3484 | 2466 |
| R-work | 0.174 | 0.188 | 0.216 |
| R-free | 0.210 | 0.233 | 0.253 |
| Number of non-hydrogen atoms | | | |
| Macromolecules | 3140 | 4704 | 4921 |
| Ligands | n/a | n/a | n/a |
| Solvent | 250 | 294 | 39 |
| Protein residues | | | |
| RMS (bonds) | 0.007 | 0.011 | 0.006 |
| RMS (angles) | 1.449 | 1.644 | 0.862 |
| Ramachandran favoured (%) | 97.2 | 97.0 | 93.8 |
| Ramachandran allowed (%) | 100.0 | 100.0 | 98.8 |
| Ramachandran outliers (%) | 0.0 | 0.0 | 1.2 |
| Rotamer outliers (%) | 0.60 | 0.40 | 0.82 |
| Clash score | 2.58 | 1.71 | 4.93 |
| Average B-factor | | | |
| Macromolecules | 22.2 | 35.7 | 69.4 |
| Ligands | n/a | n/a | n/a |
| Solvent | 31.3 | 40.8 | 60.8 |

*Statistics for the highest-resolution shell are shown in parentheses.

**Table S6b. Data collection and refinement statistics**

| | COPB2 WD-domain | KIF21A | WDR12 |
|---|---|---|---|
| PDB ID | 8D30 | 7KLJ | 6N31 |
| Wavelength (Å ) | 0.97918 | 1.54178 | 0.97918 |
| Resolution range (Å) | 50.00-2.40 (2.44-2.40) * | 50.00-1.50 (1.55-1.52) * | 76.00-2.60 (2.71-2.60) * |
| Space group | $P2_12_12_1$ | $P2_1$ | $P2_12_12_1$ |
| Unit cell (Å) | 67.1, 134.1, 144.3 | 55.3, 67.1, 79.9 | 52.7, 87.1, 155.9 |
| Total reflections | 288205 | 600649 | 65932 |
| Unique reflections | 51291 (2439) | 89050(3875) | 21949(8091) |
| Multiplicity | 5.6(4.8) | 6.7(3.9) | 3.0(3.0) |
| Completeness (%) | 99.0(97.1) | 99.3(87.1) | 96.7(95.7) |
| Mean I/sigma(I) | 21.0(1.45) | 35.8(2.98) | 7.4(1.1) |
| Wilson B-factor | 48.8 | 15.2 | 64.8 |
| R-merge | 0.080 (1.005) | 0.073(0.424) | 0.109(1.028) |
| R-meas | 0.089 (1.123) | 0.078(0.487) | 0.133(1.252) |
| R-pim | 0.037(0.486) | 0.029(0.233) | 0.046 (0.523) |
| CC1/2 | 0.998(0.496) | 0.997(0.821) | 0.993(0.476) |
| Reflections used in refinement | 51241 | 87717 | 21880 |
| Reflections used for R-free | 2516 | 1307 | 1046 |
| R-work | 0.219 | 0.170 | 0.203 |
| R-free | 0.247 | 0.185 | 0.233 |
| Number of non-hydrogen atoms | | | |
| Macromolecules | 8483 | 5004 | 4707 |
| Ligands | n/a | n/a | n/a |
| Solvent | 80 | 556 | n/a |
| Protein residues | | | |
| RMS (bonds) | 0.010 | 0.007 | 0.010 |
| RMS (angles) | 1.19 | 1.439 | 1.30 |
| Ramachandran favoured (%) | 94.6 | 95.7 | 96.1 |
| Ramachandran allowed (%) | 99.6 | 100.0 | 100.0 |
| Ramachandran outliers (%) | 0.04 | 0.0 | 0.0 |
| Rotamer outliers (%) | 1.50 | 0.92 | 3.23 |
| Clash score | 3.29 | 1.62 | 4.78 |
| Average B-factor | | | |
| Macromolecules | 61.4 | 20.1 | 49.7 |
| Ligands | n/a | n/a | n/a |
| Solvent | 50.4 | 29.8 | n/a |

*Statistics for the highest-resolution shell are shown in parentheses.

**Table S6c. Data collection and refinement statistics**

| | SEC31A | COPA | UTP15 |
|---|---|---|---|
| PDB ID | 7SUL | 6PBG | 7RUO |
| Wavelength (Å ) | 0.97918 | 1.5418 | 1.54178 |
| Resolution range (Å) | 50.00-2.40(2.44-2.40) * | 50.00-1.72(1.75-1.72) * | 42.66-1.80 (1.84-1.80) * |
| Space group | $P2_1$ | P2 | P1 |
| Unit cell (Å) | 51.2, 156.7, 85.2 | 34.6, 85.5, 47.9 | 58.3, 59.1, 61.9 |
| Total reflections | 167966 | 170533 | 125959 |
| Unique reflections | 49159(1994) | 26521(977) | 57233(3260) |
| Multiplicity | 3.4(3.0) | 6.4(4.1) | 2.2(2.2) |
| Completeness (%) | 94.2(75.0) | 93.6(69.3) | 93.7(90.0) |
| Mean I/sigma(I) | 13.1(1.46) | 21.3(2.08) | 4.7(2.0) |
| Wilson B-factor | 38.6 | 14.6 | 12.2 |
| R-merge | 0.116(0.781) | 0.088(0.472) | 0.108(0.464) |
| R-meas | 0.136(0.917) | 0.096(0.532) | 0.145(0.625) |
| R-pim | 0.069(0.473) | 0.037(0.236) | 0.096(0.415) |
| CC1/2 | 0.985(0.700) | 0.962(0.833) | 0.982(0.672) |
| Reflections used in refinement | 49130 | 25722 | 54327 |
| Reflections used for R-free | 986 | 740 | 2896 |
| R-work | 0.189 | 0.171 | 0.231 |
| R-free | 0.237 | 0.218 | 0.263 |
| Number of non-hydrogen atoms | | | |
| Macromolecules | 9956 | 2491 | 4723 |
| Ligands | n/a | n/a | n/a |
| Solvent | 69 | 220 | 292 |
| Protein residues | | | |
| RMS (bonds) | 0.010 | 0.008 | 0.007 |
| RMS (angles) | 1.20 | 1.352 | 1.490 |
| Ramachandran favoured (%) | 95.8 | 95.1 | 96.0 |
| Ramachandran allowed (%) | 100.0 | 100.0 | 100.0 |
| Ramachandran outliers (%) | 0.0 | 0.0 | 0.0 |
| Rotamer outliers (%) | 3.94 | 0.75 | 2.14 |
| Clash score | 3.56 | 2.04 | 4.7 |
| Average B-factor | | | |
| Macromolecules | 53.3 | 18.9 | 16.2 |
| Ligands | n/a | n/a | n/a |
| Solvent | 44.2 | 27.1 | 20.4 |

*Statistics for the highest-resolution shell are shown in parentheses.

**Table S6d. Data collection and refinement statistics**

| | CORO1C | RBBP7 | eIF2A |
|---|---|---|---|
| PDB ID | 7STY | 7M3X | 8DYS |
| Wavelength (Å ) | 0.97918 | 0.97918 | 0.97918 |
| Resolution range (Å) | 50.00-2.00(2.03-2.00) * | 50.00-1.46(1.49-1.46) * | 47.67-1.80(1.84-1.80) * |
| Space group | $P3_121$ | $P2_12_12_1$ | $P2_12_12_1$ |
| Unit cell (Å) | 127.2, 127.2, 59.2 | 44.7, 88.8, 97.2 | 54.3, 84.8, 99.3 |
| Total reflections | 145933 | 727132 | 326278 |
| Unique reflections | 35890(1520) | 68121(3354) | 42642(2481) |
| Multiplicity | 4.1(3.4) | 10.7(8.1) | 7.7(7.9) |
| Completeness (%) | 95.1(81.8) | 99.9(99.8) | 98.8(98.1) |
| Mean I/sigma(I) | 18.9(1.3) | 31.0(1.71) | 13.5(3.6) |
| Wilson B-factor | 40.0 | 10.4 | 23.0 |
| R-merge | 0.096(0.877) | 0.074(0.777) | 0.108(0.808) |
| R-meas | 0.110(1.021) | 0.078(0.829) | 0.117(0.864) |
| R-pim | 0.053(0.513) | 0.023(0.286) | 0.043(0.304) |
| CC1/2 | 0.990(0.507) | 0.997(0.805) | 0.995(0.832) |
| Reflections used in refinement | 34775 | 64741 | 40514 |
| Reflections used for R-free | 1097 | 3307 | 2087 |
| R-work | 0.179 | 0.170 | 0.200 |
| R-free | 0.221 | 0.193 | 0.228 |
| Number of non-hydrogen atoms | | | |
| Macromolecules | 3028 | 3085 | 3178 |
| Ligands | n/a | n/a | n/a |
| Solvent | 167 | 423 | 199 |
| Protein residues | | | |
| RMS (bonds) | 0.010 | 0.007 | 0.008 |
| RMS (angles) | 1.397 | 1.446 | 1.436 |
| Ramachandran favoured (%) | 97.4 | 98.0 | 97.1 |
| Ramachandran allowed (%) | 100.0 | 99.5 | 100.0 |
| Ramachandran outliers (%) | 0.0 | 0.5 | 0.0 |
| Rotamer outliers (%) | 0.94 | 0.88 | 0.60 |
| Clash score | 3.87 | 2.65 | 2.56 |
| Average B-factor | | | |
| Macromolecules | 43.8 | 16.2 | 28.3 |
| Ligands | n/a | n/a | n/a |
| Solvent | 44.5 | 25.9 | 33.0 |

*Statistics for the highest-resolution shell are shown in parentheses.

**Table S6e. Data collection and refinement statistics**

| | PAFAH1B1 | COPB2+OICR-6254 | WDR5+MR44397 |
|---|---|---|---|
| PDB ID | 7MT1 | 8D41 | 8T5I |
| Wavelength (Å ) | 0.97918 | 1.54178 | 0.97918 |
| Resolution range (Å) | 36.02-1.30 (1.32-1.30) * | 48.87-2.00 (2.05-2.00) * | 50.00-1.70(1.73-1.70) * |
| Space group | $P2_1$ | $P2_1$ | C2 |
| Unit cell (Å) | 36.7, 68.6, 69.5 | 57.0, 91.1, 58.7 | 101.4, 86.4, 80.8 |
| Total reflections | 464783 | 160563 | 416078 |
| Unique reflections | 82396(3866) | 38534(2656) | 72369(3548) |
| Multiplicity | 5.6(5.3) | 4.2(3.9) | 5.7(5.4) |
| Completeness (%) | 99.2(94.4) | 96.1(90.1) | 95.9(95.1) |
| Mean I/sigma(I) | 22.5(3.1) | 11.3(3.0) | 15.8(1.85) |
| Wilson B-factor | 12.6 | 10.1 | 15.9 |
| R-merge | 0.034(0.477) | 0.112(0.534) | 0.135(0.982) |
| R-meas | 0.037(0.531) | 0.128(0.617) | 0.149(1.087) |
| R-pim | 0.015(0.228) | 0.062(0.305) | 0.062(0.456) |
| CC1/2 | 1.000(0.862) | 0.993(0.779) | 0.983(0.658) |
| Reflections used in refinement | 78418 | 36560 | 70899 |
| Reflections used for R-free | 3953 | 1919 | 1467 |
| R-work | 0.177 | 0.171 | 0.197 |
| R-free | 0.197 | 0.220 | 0.232 |
| Number of non-hydrogen atoms | | | |
| Macromolecules | 2554 | 4871 | 4731 |
| Ligands | n/a | 24 | 60 |
| Solvent | 258 | 351 | 440 |
| Protein residues | | | |
| RMS (bonds) | 0.007 | 0.010 | 0.007 |
| RMS (angles) | 1.431 | 1.276 | 1.398 |
| Ramachandran favoured (%) | 95.3 | 95.6 | 95.4 |
| Ramachandran allowed (%) | 99.7 | 99.8 | 100.0 |
| Ramachandran outliers (%) | 0.3 | 0.2 | 0.0 |
| Rotamer outliers (%) | 0.0 | 0.93 | 0.95 |
| Clash score | 1.19 | 4.28 | 3.46 |
| Average B-factor | | | |
| Macromolecules | 16.5 | 20.3 | 18.7 |
| Ligands | n/a | 30.8 | 18.2 |
| Solvent | 28.7 | 25.0 | 27.5 |

*Statistics for the highest-resolution shell are shown in parentheses.

**Table S6f. Data collection and refinement statistics**

| | SETDB1+MR46747 |
|---|---|
| PDB ID | 8UWP |
| Wavelength (Å ) | 0.97918 |
| Resolution range (Å) | 50.00-1.76(1.79-1.76) * |
| Space group | $P2_12_12$ |
| Unit cell (Å) | 63.3, 141.7, 55.5 |
| Total reflections | 403142 |
| Unique reflections | 49143(2422) |
| Multiplicity | 8.2(7.7) |
| Completeness (%) | 99.0(98.8) |
| Mean I/sigma(I) | 27.6(1.83) |
| Wilson B-factor | 25.3 |
| R-merge | 0.077(0.940) |
| R-meas | 0.082(1.002) |
| R-pim | 0.028(0.341) |
| CC1/2 | 0.996(0.881) |
| Reflections used in refinement | 47896 |
| Reflections used for R-free | 1202 |
| R-work | 0.199 |
| R-free | 0.242 |
| Number of non-hydrogen atoms | |
| Macromolecules | 3529 |
| Ligands | 60 |
| Solvent | 201 |
| Protein residues | |
| RMS (bonds) | 0.009 |
| RMS (angles) | 1.363 |
| Ramachandran favoured (%) | 97.4 |
| Ramachandran allowed (%) | 100.0 |
| Ramachandran outliers (%) | 0.0 |
| Rotamer outliers (%) | 1.11 |
| Clash score | 5.21 |
| Average B-factor | |
| Macromolecules | 49.7 |
| Ligands | 34.7 |
| Solvent | 37.5 |

*Statistics for the highest-resolution shell are shown in parentheses.

**Table S11. SPR-based selectivity of seven DEL-ML compounds against six targets.**

| Compound ID | Percent binding at compound concentration of 50 µM | | | | | |
|---|---|---|---|---|---|---|
| | WDR5 | WDR12 | DCAF1 | SETDB1 | WDR91 | LRRK2 |
| MR44397 | 98 ± 4 | 23 ± 1 | 34 ± 2 | 16 ± 0.1 | 8 ± 1 | 16 ± 0.4 |
| MR40903 | 6 ± 1 | 98 ± 9 | 0 | 2 ± 0.3 | 0 | 2 ± 1 |
| OICR-6766 | 12 ± 3 | 0 | 96 ± 2 | 3 ± 0.1 | 0 | 7 ± 1 |
| MR46747 | 1 ± 0.1 | 0 | 0 | 83 ± 3 | 0 | 0 |
| MR45279 | 0 | 0 | 0 | 0 | 77 ± 8 | 2 ± 1 |
| DR02034* | 0 | 1 ± 0.1 | 0 | 0 | 0 | 15 ± 1 |
| DR02380* | 0 | 0 | 0 | 0 | 2 ± 1 | 12 ± 4 |

*Visible precipitation in stock solution; please see Table S8 for solubility data.

**Figure S1**

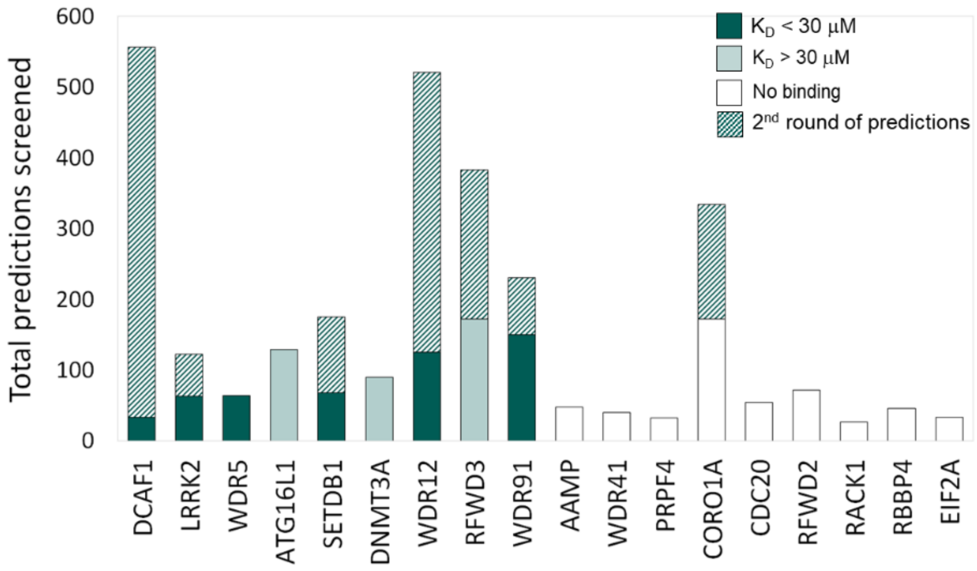


**Fig. S1 | The total number of predicted ligands tested**. The SMILES are listed in **Table S8**.

**Figure S2**

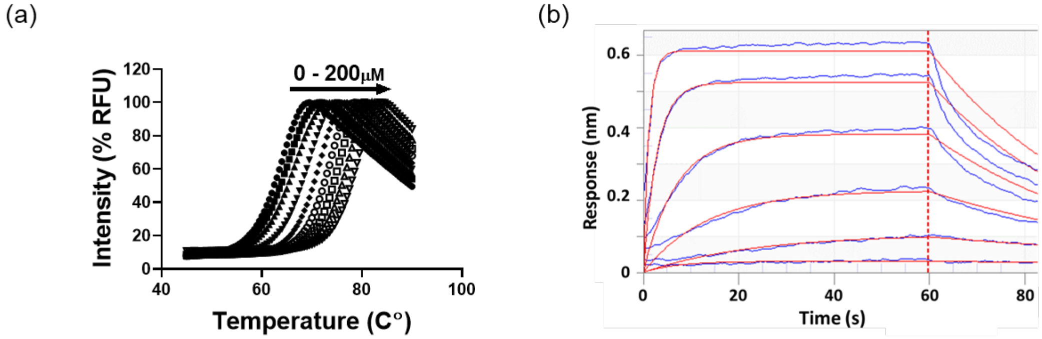


**Fig. S2 | DSF and BLI data for WDR5. (a)** Thermal stability of WDR5 was assessed in the presence and absence of MR44397 using differential scanning fluorimetry (DSF). Effect of MR44397 binding on the stability of WDR5 was measured by DSF at 0.1 mg/mL WDR5 in the presence of 0-200 µM of MR44397. **(b)** BLI analysis of the binding of MR44397 to WDR5. A representative sensorgram (blue) is globally fitted (red) with a $K_D$ value of 497 ± 32 nM, $k_{on}$: 0.6 ± 0.1 × $10^{+5}$ $M^{-1}$ $s^{-1}$; $k_{off}$: 2.8 ± 0.1 × $10^{-2}$ $s^{-1}$. All experiments were performed in triplicate.

Figure S3

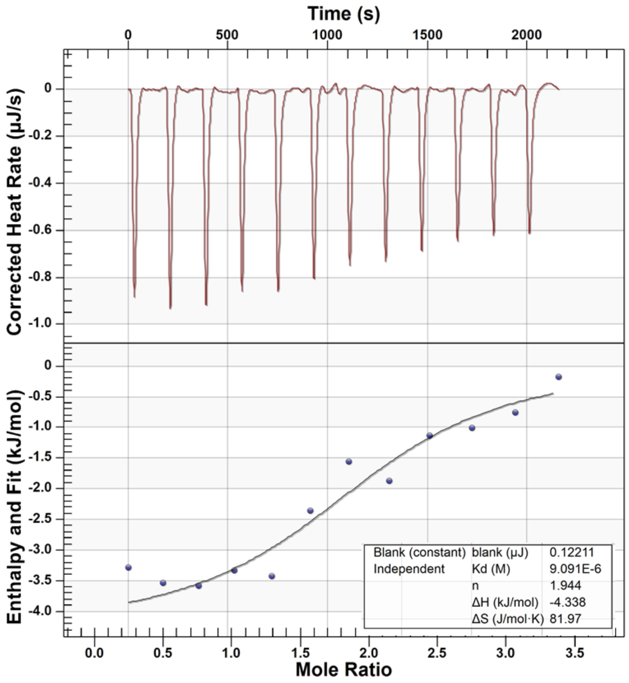


**Fig. S3 | ITC for** Z1391232269/**compound 3c in Ref 14.**

Figure S4

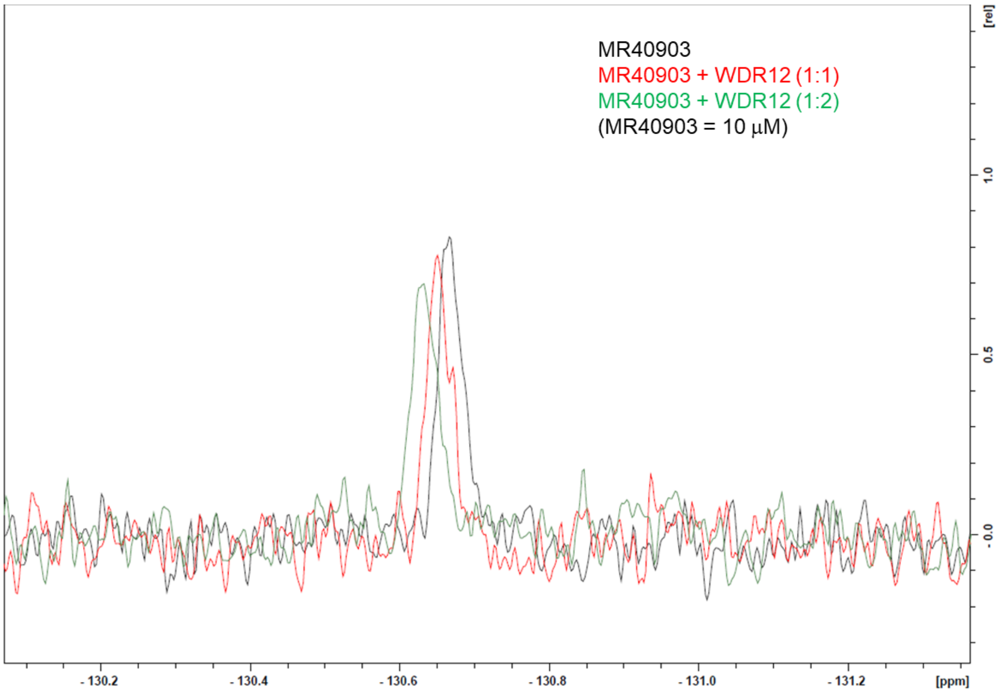


**Fig. S4 |** Ligand observed $^{19}$F-NMR for primary hit MR40903 binding to WDR12.

Figure S5

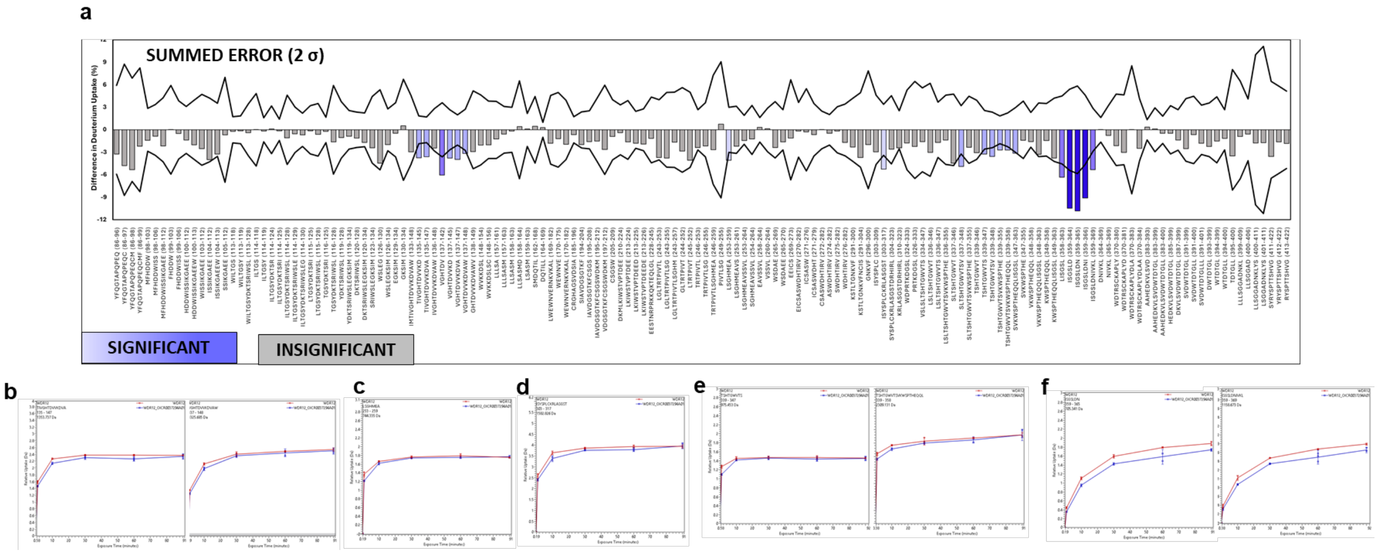


**Figure S5. HDX-MS Kinetic Plots of WDR12 peptides spanning residues** (B) 135-148, (C) 253-259, (D) 303-317, (E) 339-358, and (F) 359-369. The deuterium uptake of WDR12 (red) and WDR12+MR44915 1:20 (blue) was tracked across 1, 10, 30, 60, and 90 mins. Divergence in the red and blue lines indicate perturbations in conformational dynamics due to binding MR44915. Complexation caused B-E to experience decreased deuterium uptake early in the time course whereas residues 359-369 (in F) had persistently reduced uptake.

**Fig S6**. Examples of protein capture QC used for DEL selections. I = input; FT = flow-through; R = recovery

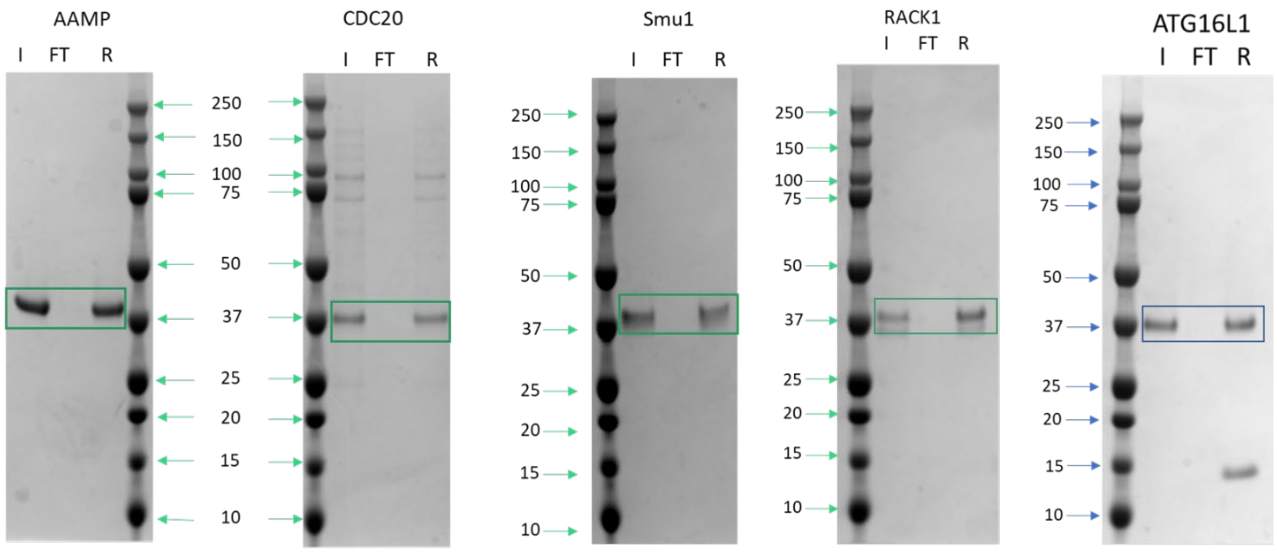

Figure S7. HPLC-UV/MS for compounds in Table 1 and Figs. 3-8.

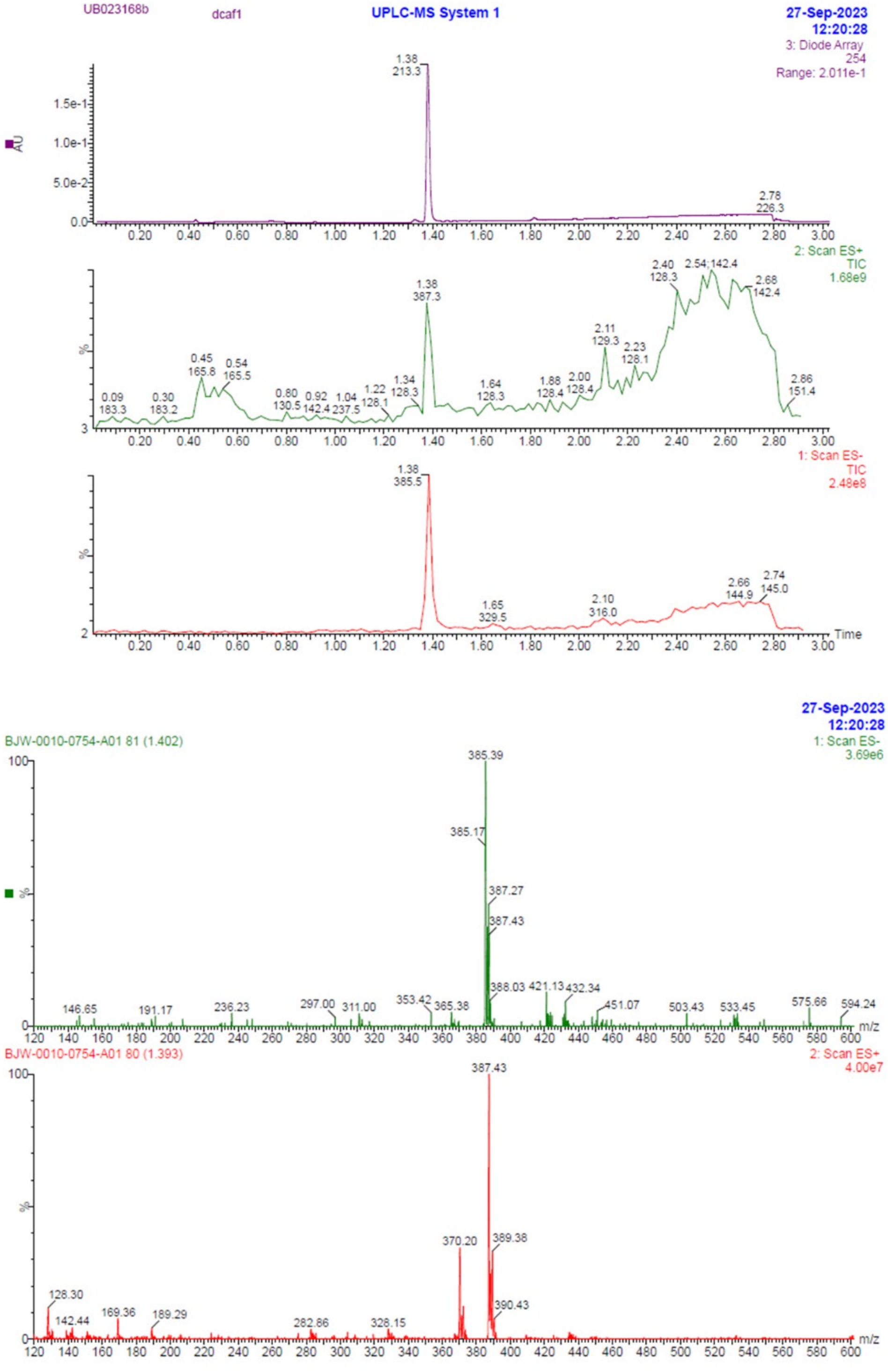

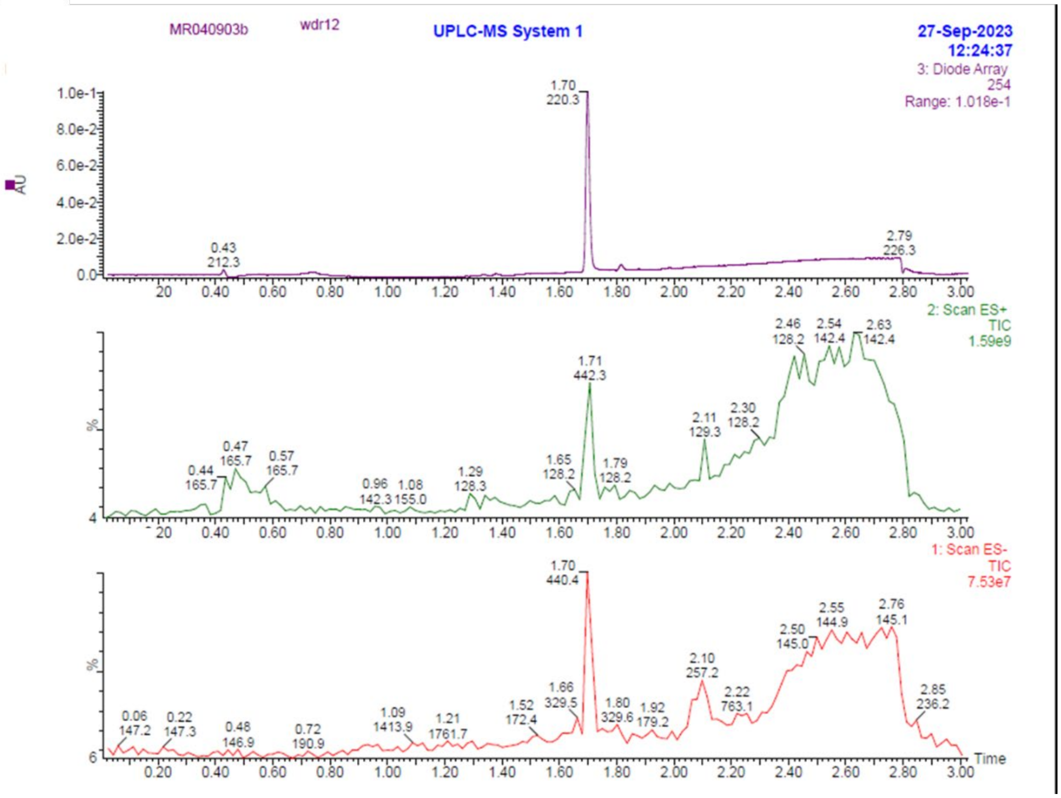
MR040903b
wdr12
UPLC-MS System 1
27-Sep-2023
12:24:37
3: Diode Array
254
Range: 1.018e-1
AU
1.70
220.3
0.43
212.3
2.79
226.3
2: Scan ES+
TIC
1.59e9
%
0.44 165.7
0.47 165.7
0.57 165.7
0.96 142.3
1.08 155.0
1.29 128.3
1.65 128.2
1.71 442.3
1.79 128.2
2.11 129.3
2.30 128.2
2.46 128.2
2.54 142.4
2.63 142.4
1: Scan ES-
TIC
7.53e7
0.06 147.2
0.22 147.3
0.48 146.9
0.72 190.9
1.09 1413.9
1.21 1761.7
1.52 172.4
1.66 329.5
1.70 440.4
1.80 329.6
1.92 179.2
2.10 257.2
2.22 763.1
2.50 145.0
2.55 144.9
2.76 145.1
2.85 236.2
Time

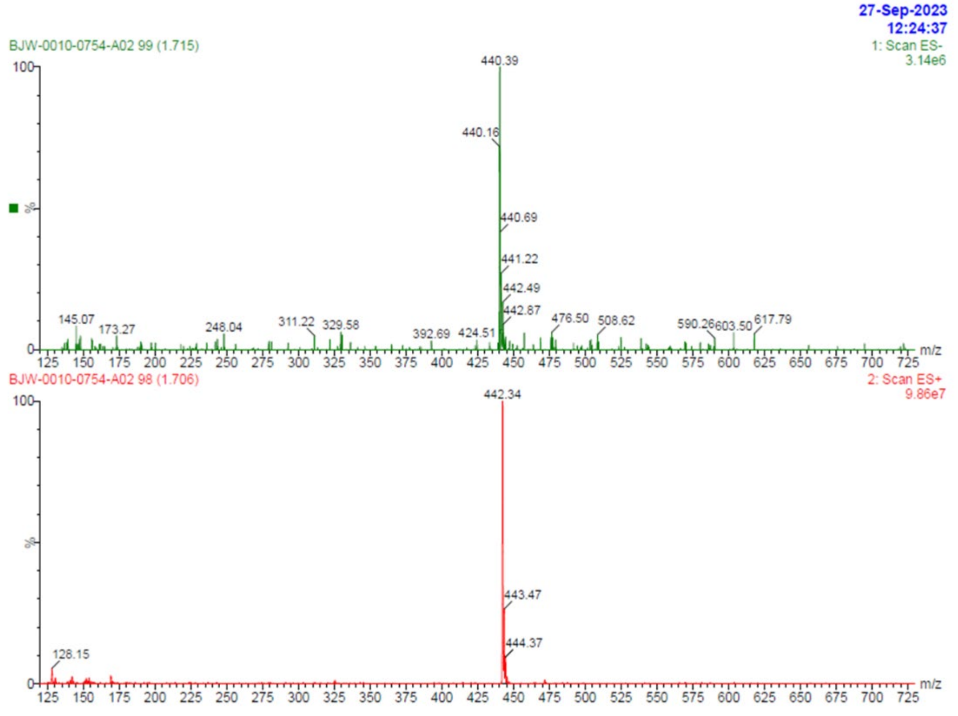
27-Sep-2023
12:24:37
BJW-0010-0754-A02 99 (1.715)
1: Scan ES-
3.14e6
%
440.39
440.16
440.69
441.22
442.49
442.87
145.07
173.27
248.04
311.22
329.58
392.69
424.51
476.50
508.62
590.26
603.50
617.79
m/z
BJW-0010-0754-A02 98 (1.706)
2: Scan ES+
9.86e7
442.34
443.47
444.37
128.15
m/z

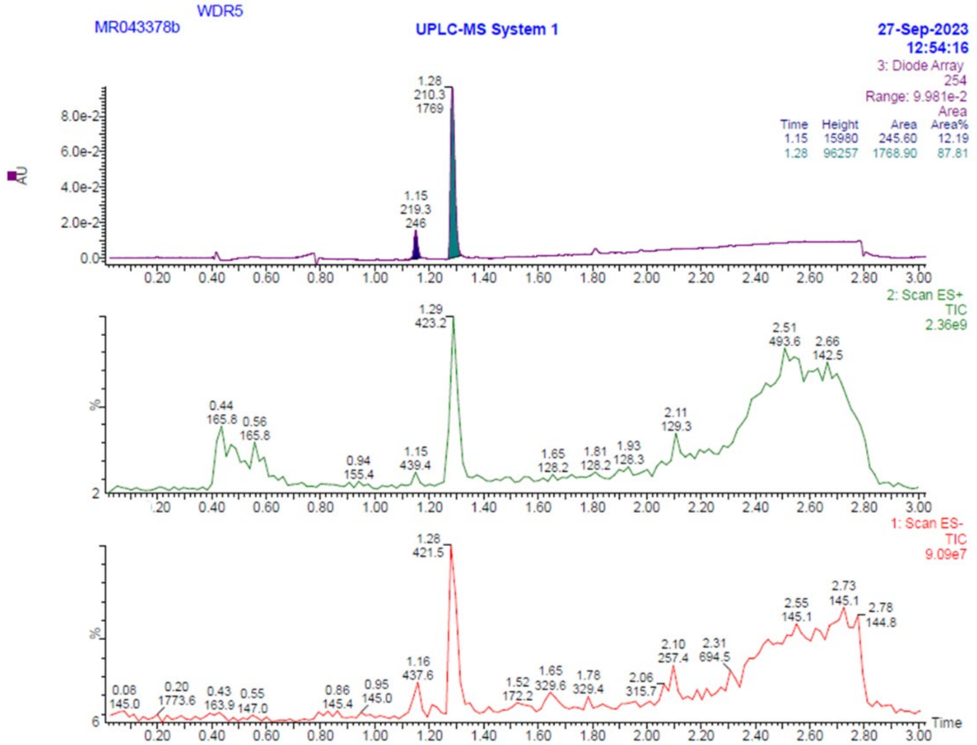
MR043378b
WDR5
UPLC-MS System 1
27-Sep-2023
12:54:16
3: Diode Array
254
Range: 9.981e-2
Area
Time Height Area Area%
1.15 15980 245.60 12.19
1.28 96257 1768.90 87.81
1.28
210.3
1769
1.15
219.3
246
AU
2: Scan ES+
TIC
2.36e9
1.29
423.2
1: Scan ES-
TIC
9.09e7
1.28
421.5
Time

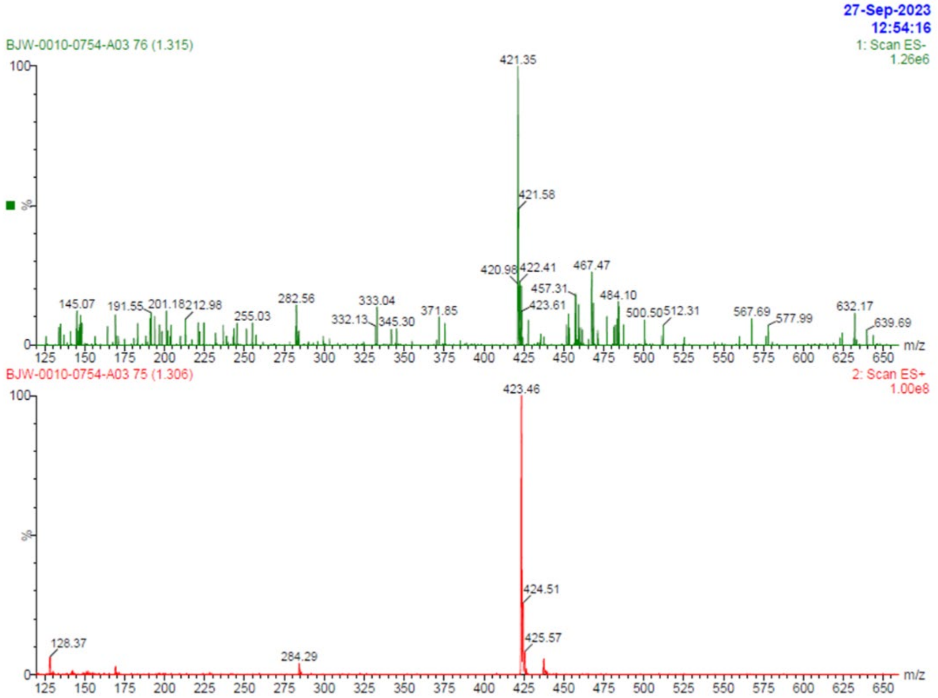
27-Sep-2023
12:54:16
BJW-0010-0754-A03 76 (1.315)
1: Scan ES-
1.26e6
421.35
421.58
420.98
422.41
467.47
457.31
423.61
484.10
m/z
BJW-0010-0754-A03 75 (1.306)
2: Scan ES+
1.00e8
423.46
424.51
425.57
128.37
284.29
m/z

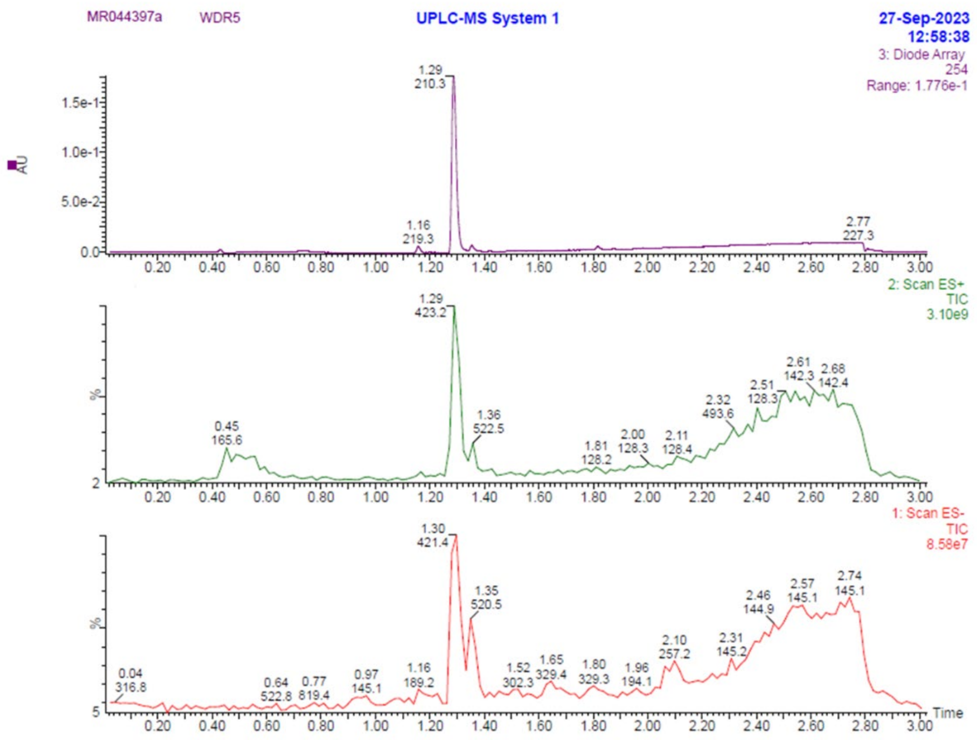
MR044397a
WDR5
UPLC-MS System 1
27-Sep-2023
12:58:38
3: Diode Array
254
Range: 1.776e-1
AU
1.5e-1
1.0e-1
5.0e-2
0.0
1.29
210.3
1.16
219.3
2.77
227.3
2: Scan ES+
TIC
3.10e9
1.29
423.2
0.45
165.6
1.36
522.5
1.81
128.2
2.00
128.3
2.11
128.4
2.32
493.6
2.51
128.3
2.61
142.3
2.68
142.4
1: Scan ES-
TIC
8.58e7
1.30
421.4
1.35
520.5
0.04
316.8
0.64
522.8
0.77
819.4
0.97
145.1
1.16
189.2
1.52
302.3
1.65
329.4
1.80
329.3
1.96
194.1
2.10
257.2
2.31
145.2
2.46
144.9
2.57
145.1
2.74
145.1
Time

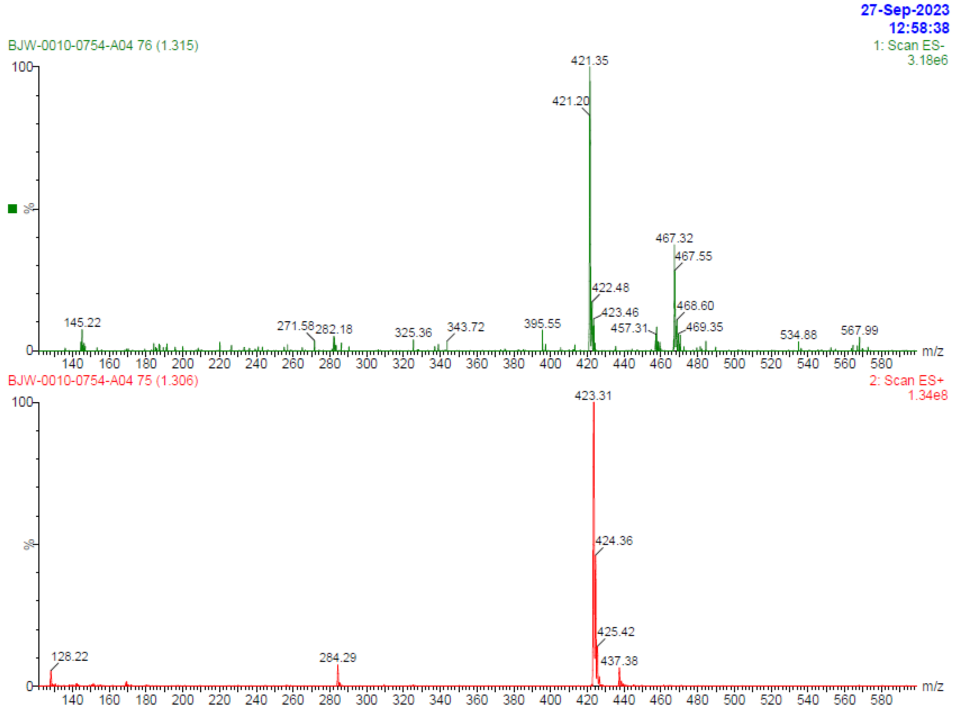
27-Sep-2023
12:58:38
BJW-0010-0754-A04 76 (1.315)
1: Scan ES-
3.18e6
421.35
421.20
467.32
467.55
422.48
423.46
468.60
457.31
469.35
145.22
271.58
282.18
325.36
343.72
395.55
534.88
567.99
m/z
BJW-0010-0754-A04 75 (1.306)
2: Scan ES+
1.34e8
423.31
424.36
425.42
437.38
128.22
284.29
m/z

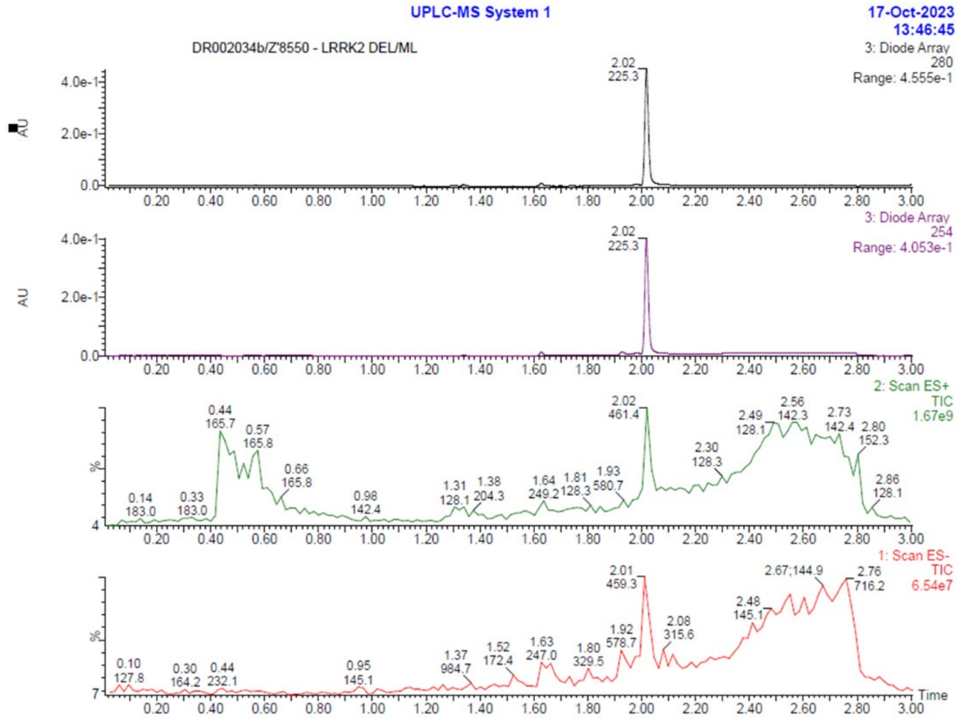
UPLC-MS System 1
17-Oct-2023
13:46:45
DR002034b/Z'8550 - LRRK2 DEL/ML
3: Diode Array
280
Range: 4.555e-1
2.02
225.3
3: Diode Array
254
Range: 4.053e-1
2: Scan ES+
TIC
1.67e9
1: Scan ES-
TIC
6.54e7
Time

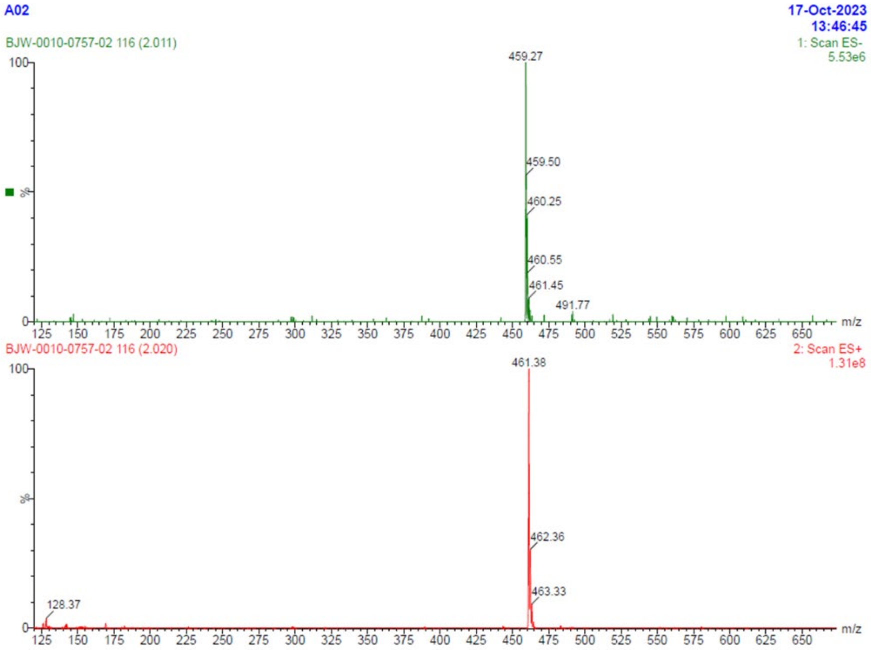
A02
17-Oct-2023
13:46:45
BJW-0010-0757-02 116 (2.011)
1: Scan ES-
5.53e6
459.27
459.50
460.25
460.55
461.45
491.77
m/z
BJW-0010-0757-02 116 (2.020)
2: Scan ES+
1.31e8
461.38
462.36
463.33
128.37
m/z

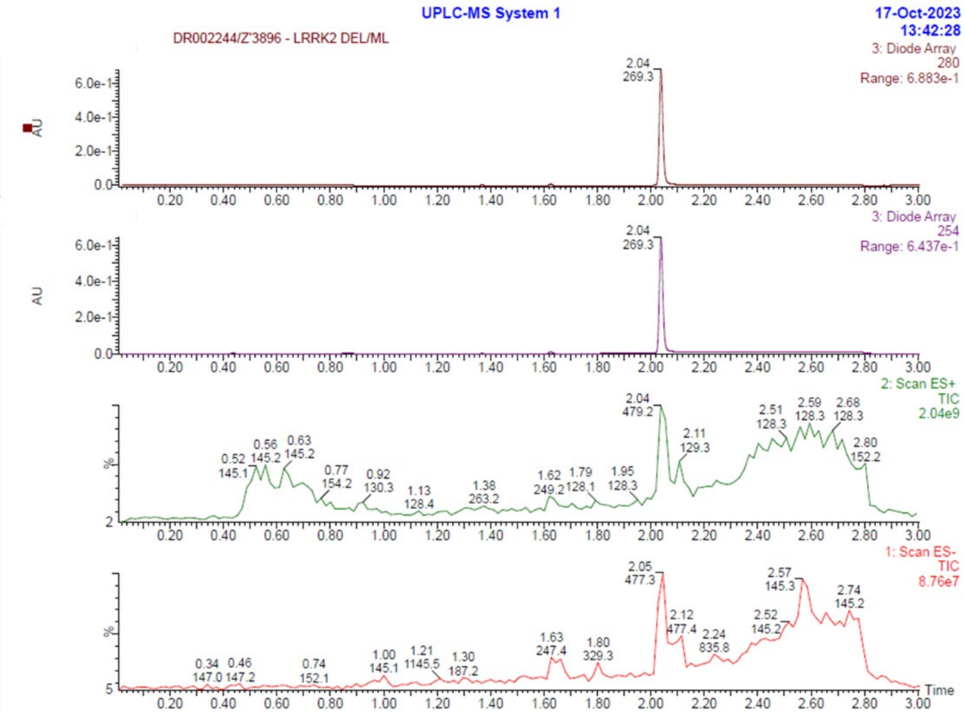
UPLC-MS System 1
17-Oct-2023
13:42:28
DR002244/Z'3896 - LRRK2 DEL/ML
3: Diode Array
280
Range: 6.883e-1
2.04
269.3
AU
3: Diode Array
254
Range: 6.437e-1
2.04
269.3
AU
2: Scan ES+
TIC
2.04e9
2.04
479.2
0.52 145.1
0.56 145.2
0.63 145.2
0.77 154.2
0.92 130.3
1.13 128.4
1.38 263.2
1.62 249.2
1.79 128.1
1.95 128.3
2.11 129.3
2.51 128.3
2.59 128.3
2.68 128.3
2.80 152.2
%
1: Scan ES-
TIC
8.76e7
2.05
477.3
0.34 147.0
0.46 147.2
0.74 152.1
1.00 145.1
1.21 1145.5
1.30 187.2
1.63 247.4
1.80 329.3
2.12 477.4
2.24 835.8
2.52 145.2
2.57 145.3
2.74 145.2
%
Time

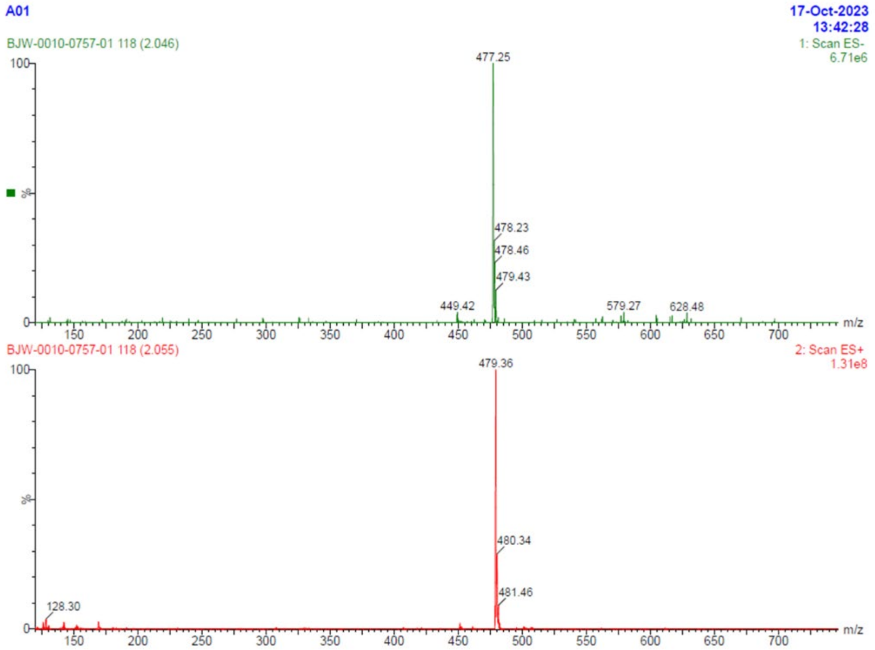
A01
17-Oct-2023
13:42:28
BJW-0010-0757-01 118 (2.046)
1: Scan ES-
6.71e6
477.25
478.23
478.46
479.43
449.42
579.27
628.48
%
m/z
BJW-0010-0757-01 118 (2.055)
2: Scan ES+
1.31e8
479.36
480.34
481.46
128.30
%
m/z

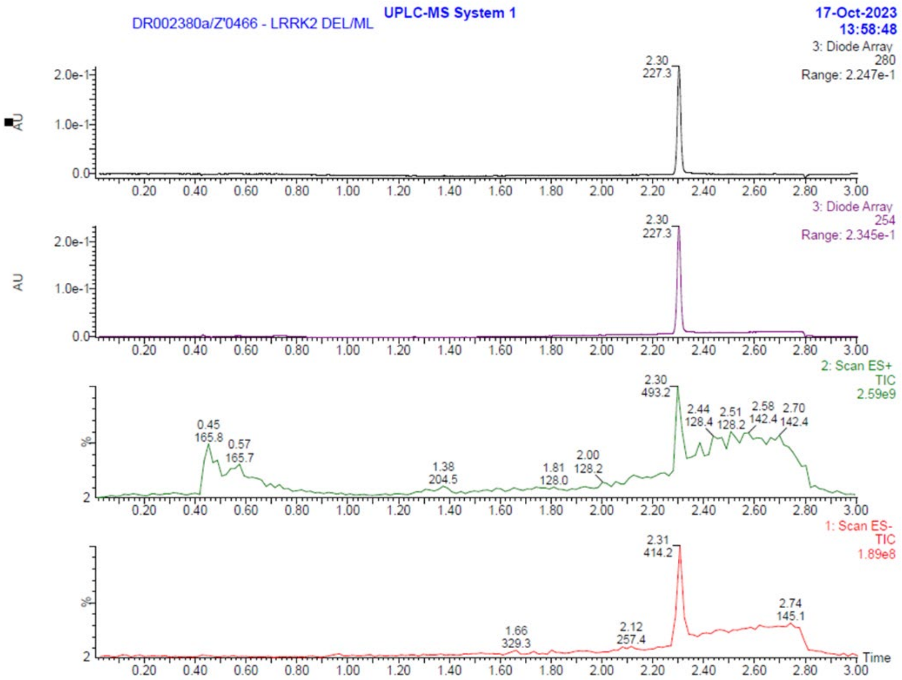
DR002380a/Z'0466 - LRRK2 DEL/ML
UPLC-MS System 1
17-Oct-2023
13:58:48
3: Diode Array
280
Range: 2.247e-1
2.30
227.3
3: Diode Array
254
Range: 2.345e-1
2.30
227.3
2: Scan ES+
TIC
2.59e9
2.30
493.2
0.45
165.8
0.57
165.7
1.38
204.5
1.81
128.0
2.00
128.2
2.44
128.4
2.51
128.2
2.58
142.4
2.70
142.4
1: Scan ES-
TIC
1.89e8
2.31
414.2
1.66
329.3
2.12
257.4
2.74
145.1
AU
%
Time

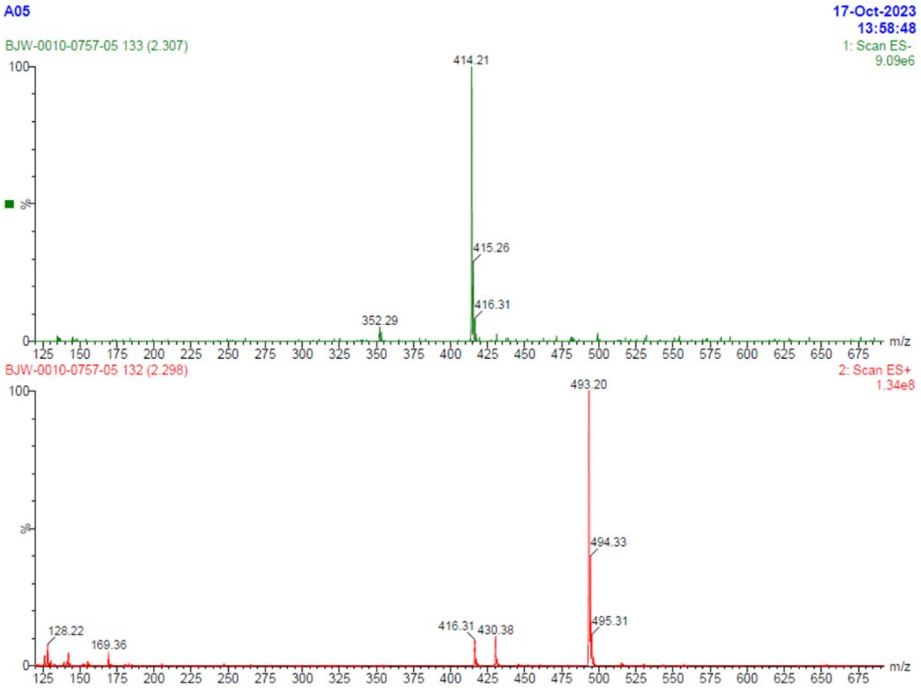
A05
17-Oct-2023
13:58:48
BJW-0010-0757-05 133 (2.307)
1: Scan ES-
9.09e6
414.21
415.26
416.31
352.29
m/z
BJW-0010-0757-05 132 (2.298)
2: Scan ES+
1.34e8
493.20
494.33
495.31
416.31
430.38
128.22
169.36
m/z
%
